# Increased sensitivity of SARS-CoV-2 to type III interferon in human intestinal epithelial cells

**DOI:** 10.1101/2021.06.14.448464

**Authors:** Carmon Kee, Camila Metz-Zumaran, Patricio Doldan, Cuncai Guo, Megan L. Stanifer, Steeve Boulant

## Abstract

The coronavirus SARS-CoV-2 caused the COVID-19 global pandemic leading to 3.5 million deaths worldwide as of June 2021. The human intestine was found to be a major viral target which could have a strong impact on virus spread and pathogenesis since it is one of the largest organs. While type I interferons (IFNs) are key cytokines acting against systemic virus spread, in the human intestine type III IFNs play a major role by restricting virus infection and dissemination without disturbing homeostasis. Recent studies showed that both type I and III IFNs can inhibit SARS-CoV-2 infection, but it is not clear if one IFN controls SARS-CoV-2 infection of the human intestine better or with a faster kinetics. In this study, we could show that both type I and III IFNs possess antiviral activity against SARS-CoV-2 in human intestinal epithelial cells (hIECs), however type III IFN is more potent. Shorter type III IFN pretreatment times and lower concentrations were required to efficiently reduce virus load when compared to type I IFNs. Moreover, type III IFNs significantly inhibited SARS-CoV-2 even 4 hours post-infection and induced a long-lasting antiviral effect in hIECs. Importantly, the sensitivity of SARS-CoV-2 to type III IFNs was virus-specific since type III IFN did not control VSV infection as efficiently. Together these results suggest that type III IFNs have a higher potential for IFN-based treatment of SARS-CoV-2 intestinal infection as compared to type I IFNs.

## Introduction

Since the end of 2019 we have witnessed a global pandemic due to the emergence of the severe-acute-respiratory-syndrome-related coronavirus-2 (SARS-CoV-2) (Lu et al., 2020). Coronaviruses are enveloped, single-stranded positive-sense RNA viruses that can infect most animal species and generally cause common colds in humans (Haake et al., 2020). However, in the past 19 years coronaviruses have been involved in zoonotic events giving rise to highly pathogenic human viruses (e.g. MERS and SARS-CoV-1). The latest, SARS-CoV-2, is responsible for the coronavirus-associated acute respiratory disease or coronavirus disease 19 (COVID-19) and as of June 2021 has caused more than 150 million infections and 3.5 million deaths worldwide (World Health Organization 2021). The virus and its associated disease have caused a worldwide medical and economic impact and united efforts are necessary to find adequate antiviral treatments.

The spread of the virus among humans is thought to occur through respiratory droplets. Thereby, SARS-CoV-2 primarily targets cells of the lung epithelium, causing respiratory symptoms ranging from cough and shortness of breath to severe lung injury (Huang et al., 2020; Zhu et al., 2020). Interestingly, a significant number of patients with COVID-19 report gastrointestinal symptoms including diarrhea, nausea or vomiting, that can either precede or follow respiratory symptoms (Guan et al., 2020; Huang et al., 2020; L. Lin et al., 2020). Moreover, multiple reports detected viral RNA in feces, and stool specimens of infected patients remained positive even after a negative oropharyngeal swab test (Wölfel et al., 2020; Y. Wu et al., 2020; Xiao et al., 2020; Xing et al., 2020; Xu et al., 2020). Direct evidence for viral infection of gut tissue was also shown through evaluation of endoscopic samples (Xiao et al., 2020). Importantly, within the past months the understanding of intestinal SARS-CoV-2 infection and pathogenesis was improved with *in vitro* studies. Several studies demonstrated that immortalized human intestinal cells and primary human mini-gut organoids supported SARS-CoV-2 infection (Lamers et al., 2020; Stanifer, Guo, et al., 2020; Triana et al., 2021; Zang et al., 2020). Altogether, the medical reports and scientific data clearly show that SARS-CoV-2 is not restricted to the respiratory tract but can infect the human intestinal epithelium. However, it remains unknown to which extent the enteric phase is important for viremia, pathogenesis, and transmission.

Most cells in the body respond to viral infection by generating both a proinflammatory and an interferon (IFN) response. Secreted IFNs bind to IFN receptors inducing a signaling cascade leading to the transcription of hundreds of interferon stimulated genes (ISGs) which in turn create an antiviral state (Stanifer et al., 2019). Almost all cells in the body produce and respond to type I IFN to mount their antiviral response. Interestingly, the response of both lung and intestinal epithelial cells to viral infection also strongly depends on an additional interferon, the type III IFN. This tropism for the type III IFN action is due to the fact that the type III IFN receptor is mainly expressed on epithelial cells, placing it as a unique antiviral strategy for epithelium and mucosal surfaces. Several studies have demonstrated that during enteric virus infection, the type I IFNs are essential to protect against systemic spread while the type III IFNs maintain epithelial balance protecting intestinal epithelial cells while limiting an exacerbated immune response (Baldridge et al., 2015; Dionne et al., 2011; Mahlakõiv et al., 2015; Nice et al., 2015; Pott et al., 2011; Pott & Stockinger, 2017; Weber et al., 2014) and this observation also stands true for lung respiratory epithelium (Klinkhammer et al., 2018; Luker et al., 2003). Importantly, while human intestinal epithelial cells can respond to both type I and III IFNs to control virus replication and spread when exogenously provided (Pervolaraki et al., 2017, 2018; Stanifer, Guo, et al., 2020), evidence suggests that the endogenous type III IFN are key to protect against virus infection (Stanifer, Kee, et al., 2020). The function of type I and III IFNs in the respiratory tract appears to be dependent on the location of the infection. Within the upper respiratory tract, type III IFNs are key to control influenza viruses (Klinkhammer et al., 2018) while in the lower airway epithelium, both type I and type III IFNs have redundant functions in controlling viral infection (Crotta et al., 2013).

Infection of human lung epithelial cells by SARS-CoV-2 was reported to induce a typical IFN response shown by the upregulation of the IFN themselves as well as the induction of ISGs (Desai et al., 2020; Katsura et al., 2020). Interestingly, other reports described a limited to absent IFN response upon SARS-CoV-2 infection of lung epithelial cells (Blanco-Melo et al., 2020; Chu et al., 2021; Shuai et al., 2020; Vanderheiden et al., 2020) which could be the results of the mechanisms developed by SARS-CoV-2 to interfere with both the production of IFN and its downstream signaling (Meffre & Iwasaki, 2020; Triana et al., 2021). The impact of SARS-CoV-2 on human intestinal epithelial cells is much less characterized compared to the lung epithelial cells, however, there is clear evidence showing that a typical type I and type III IFN intrinsic innate immune response is generated upon infection (Chu et al., 2021; Lamers et al., 2020; Stanifer, Kee, et al., 2020).

Treatment of both lung and intestinal epithelial cells with type I IFN was reported to partially protect against SARS-CoV-2 infection (Felgenhauer et al., 2020; Katsura et al., 2020; Park & Iwasaki, 2020; Rebendenne et al., 2021; Shuai et al., 2020; Vanderheiden et al., 2020). Similarly pretreatment of epithelial cells with type III IFN was also reported to interfere with SARS-CoV-2 infection/replication (Busnadiego et al., 2020; Felgenhauer et al., 2020; Stanifer, Kee, et al., 2020; Vanderheiden et al., 2020). Interestingly, SARS-CoV-2 infection of human intestinal organoids, revealed that upon infection, primary human intestinal epithelial cells favor the production and secretion of type III IFN (Stanifer, Kee, et al., 2020). Additionally, deletion of either the type I IFN receptor or the type III IFN receptor from human intestinal epithelial cells revealed that the type III IFNs rather than type I IFNs had a predominant role in controlling SARS-CoV-2 infection (Stanifer, Kee, et al., 2020). The molecular origins for this more important function of type III IFN in protecting against SARS-CoV-2 in human intestinal epithelial cells remains unclear

While it is now clear that the human intestinal epithelium mounts an IFN-dependent response upon SARS-CoV-2 infection, there is very little data about the kinetics of IFN-mediated protection. In this study we investigate in detail how type I and III IFNs establish their antiviral program against SARS-CoV-2 infection in human intestinal epithelial cells. Our results showed that endogenous type I IFNs play a minor role in inhibiting SARS-CoV-2 infection, however over time endogenous type III IFNs play an essential role in controlling virus spread. While both type I and III IFNs induced an antiviral activity against SARS-CoV-2 at high concentrations, low concentrations and shorter pre-treatment of type III IFNs were sufficient to inhibit virus infection. The sensitivity of SARS-CoV-2 to type III IFNs is virus specific, since pretreatment of human intestinal epithelial cells with the same IFN concentrations and for the same amount of time did not have the same deleterious effect on VSV infection as on SARS-CoV-2 infection. Importantly, type III IFN were able to have a long-lasting effect, eliciting an antiviral state even 72h post treatment indicating that using type III IFNs as antiviral measures could have strong potential to clear infection and *de novo* virus shedding from the human intestine.

## Results

### Endogenous type III IFNs control SARS-CoV-2 replication and spread in human intestinal epithelial cells

As growing evidence supports that the gastrointestinal tract can be infected by SARS-CoV-2, studying the antiviral immune response of human intestinal epithelial cells (hIECs) is essential to understand COVID-19 pathogenesis. We previously reported that type III IFNs are critical to control SARS-CoV-2 infection in hIECs (Stanifer, Kee, et al., 2020). This was further supported by the observation that knocking out the type III IFN receptor leads to a greater increase of SARS-CoV-2 infection, replication, and *de novo* virus production compared to the knockout of the type I IFN receptor (Stanifer, Kee, et al., 2020). These previous findings were performed at a single time point post-interferon treatment and only gave a static view of type III IFNs role in controlling SARS-CoV-2 infection. To address how type I and type III IFNs play a role in confining virus replication and spread over time in hIECs, we infected the colon carcinoma cells T84 wild type or T84 cells depleted of the type I IFN receptor (IFNAR^−/−^), the type III IFN receptor (IFNLR^−/−^) or both IFN receptors (double knockout [dKO]) with SARS-CoV-2 at a MOI of 0.04 (as determined in Vero cells). At different times post infection, cells were fixed and immunostained using an antibody targeting the viral nucleocapsid protein (Figure 1A). Results show that at early time post infection (i.e. 4 and 8 hours post-infection (hpi)), cells depleted of the type I IFN receptor (IFNAR^−/−^) and cells depleted of both IFN receptors (dKO) were significantly more infected compared to WT T84 cells and cells depleted of the type III IFN receptor (Figure 1B). On the contrary, at later time post-infection (i.e. 12 and 24 hpi), cells depleted of the type III IFN receptor were found to be more infectable compared to cells depleted of the type I IFN receptor (Figure 1B). These findings confirm that both the type I and type III IFN systems are important to control SARS-CoV-2 infection in hIECs and suggest that the type I IFN pathway might play a greater protective role at earlier time point compared to the type III IFN pathway.

**Figure 1:**
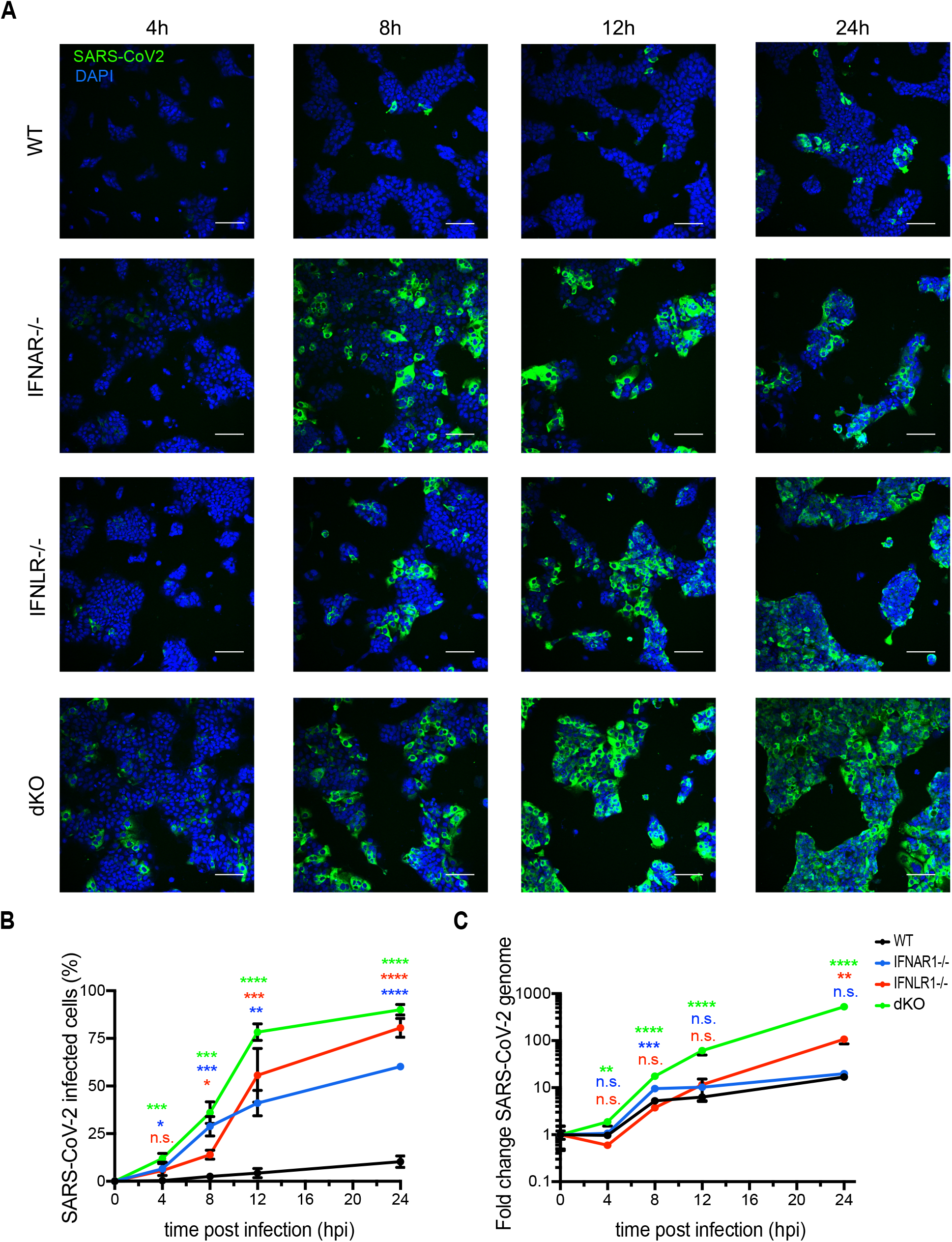
Kinetics of SARS-CoV-2 infection in human intestinal epithelial cells. (A-C) Wild type T84 cells and T84 cells depleted of either the type I IFN receptor (IFNAR^−/−^), the type III IFN receptor (IFNLR^−/−^) or depleted of both IFN receptors (dKO) were infected with SARS-CoV-2 at an MOI of 0.04 (as determined in Vero cells). At 4, 8, 12 and 24 hpi, cells were harvested to assess virus infection and replication. (A) Indirect immunofluorescence was performed against the viral nucleocapsid protein (green). Nuclei were stained with DAPI (blue). Representative images are shown. Scale bar=100 μm. (B) Same as (A) except the percentage of SARS-CoV-2-positive cells was quantified. (C) RNA was harvested, and q-RT-PCR was used to evaluate the replication of the copy number of the SARS-CoV-2 genome. Data are normalized to input. (B-C) Error bars indicate standard deviation. n = 3 biological replicates. n.s=not significant. P<0.05 *, P<0.01 **, P<0.001 ***, P <0.0001 ****, as determined by ordinary one-way ANOVA with Dunnett’s multiple comparison test using WT T84 cells as reference. The color of significance stars represents the cell line that is compared to WT T84 (green for dKO, red for IFNLR^−/−^ and blue for IFNAR^−/−^).

To address if virus replication was also increased in cells depleted of interferon receptors, T84 wildtype and interferon receptor knock-out cells were infected with SARS-CoV-2 and virus replication was monitored over time by q-RT-PCR. In agreement with our previous findings (Stanifer, Kee, et al., 2020), results show cells depleted of both IFN receptors supported a greater SARS-CoV-2 replication (Figure 1C). Importantly, and similar to the number of infected cells (Figure 1B), IFNLR^−/−^ cells showed higher levels of viral copy number compared to WT T84 and IFNAR^−/−^ cells at late times post-infection (Figure 1C). Interestingly, at late time post-infection (i.e. 24 hpi), IFNAR^−/−^ cells show a similar level of SARS-CoV-2 replication compared to WT cells (Figure 1C). Our results confirm that the absence of IFN signaling favors SARS-CoV-2 infection, replication and spread. Cells depleted of the type I IFN receptor appear to be more infectable and support more viral replication at early time post-infection compared to cells depleted of the type III IFN receptor. Conversely, the type III IFN pathways appear to play a more fundamental protective function against SARS-CoV-2 at later times post-infection. Together, these findings suggest that both IFNs act on hIECs with distinct kinetics and likely interfere with different steps of the SARS-CoV-2 lifecycle (e.g entry/replication in primary infection *vs.* spread and secondary infection).

### IFNs inhibit SARS-CoV-2 infection in a concentration dependent manner

Our data shows that endogenous type III IFN-mediated signaling is critical to control SARS-CoV-2 infection in hIECs, while endogenous type I IFNs may play a less important protective role limited to early times post-infection. However, we and other have previously reported that both type I and type III IFNs mediate an antiviral protection and that both IFNs are able to restrict SARS-CoV-2 virus replication (Felgenhauer et al., 2020; Pervolaraki et al., 2018; Stanifer, Kee, et al., 2020). To directly test the efficiency of IFNs in controlling SARS-CoV-2 infection and to address whether type I and III IFNs have a different kinetics and efficiency of antiviral activity, WT T84 cells were either mock-treated or pretreated with increasing concentrations of type I (IFNβ1) or type III (IFNλ1/2/3) IFNs for 12 or 24 hours prior to infection with SARS-CoV-2 at a MOI of 0.04 (as determined in Vero cells) (Figure 2A). At 24 hpi, cells were analyzed for virus infection by either immunofluorescence for the SARS-CoV-2 nucleoprotein or by q-RT-PCR against the virus genome. Immunofluorescence staining revealed that both type I and type III IFNs could reduce SARS-CoV-2 infection in a dose dependent manner (Figure 2B-C) and quantification of the number of infected cells revealed that both IFNs significantly impaired SARS-CoV-2 infection in a dose- and time-dependent manner (Figure 2C). Intriguingly, we noticed that for cells treated with low concentrations of type I IFNs for 12 hours, we had a greater inhibition of SARS-CoV-2 infection as compared to a higher amount of IFN. While the molecular reason for this inverted dose dependent response (at low concentrations) are not known, it possibly because at the low concentrations of IFNs an antiviral state can be induced while activation of the negative regulatory feedback loop of type I IFNs is poorly activated, resulting in a greater antiviral state compared to higher concentrations. Interestingly, high doses of type I IFN were necessary to reduce SARS-CoV-2 infection to 10% of the cells (>2000 U/mL) (Figure 2B-C). On the contrary, type III IFNs were able to restrict virus infection even at the lowest concentration (0.003 ng/mL) to below 5% infected cells (Figure 2B-C). Concomitantly to the observed reduction in the number of SARS-CoV-2 infected cells upon IFN treatment (Figure 2C), both type I and type III IFNs lead to a dose and time dependent decrease in viral replication as monitored by evaluating the relative increase of SARS-CoV-2 genome copy number over a 24 hrs infection (Figure 2D).

**Figure 2:**
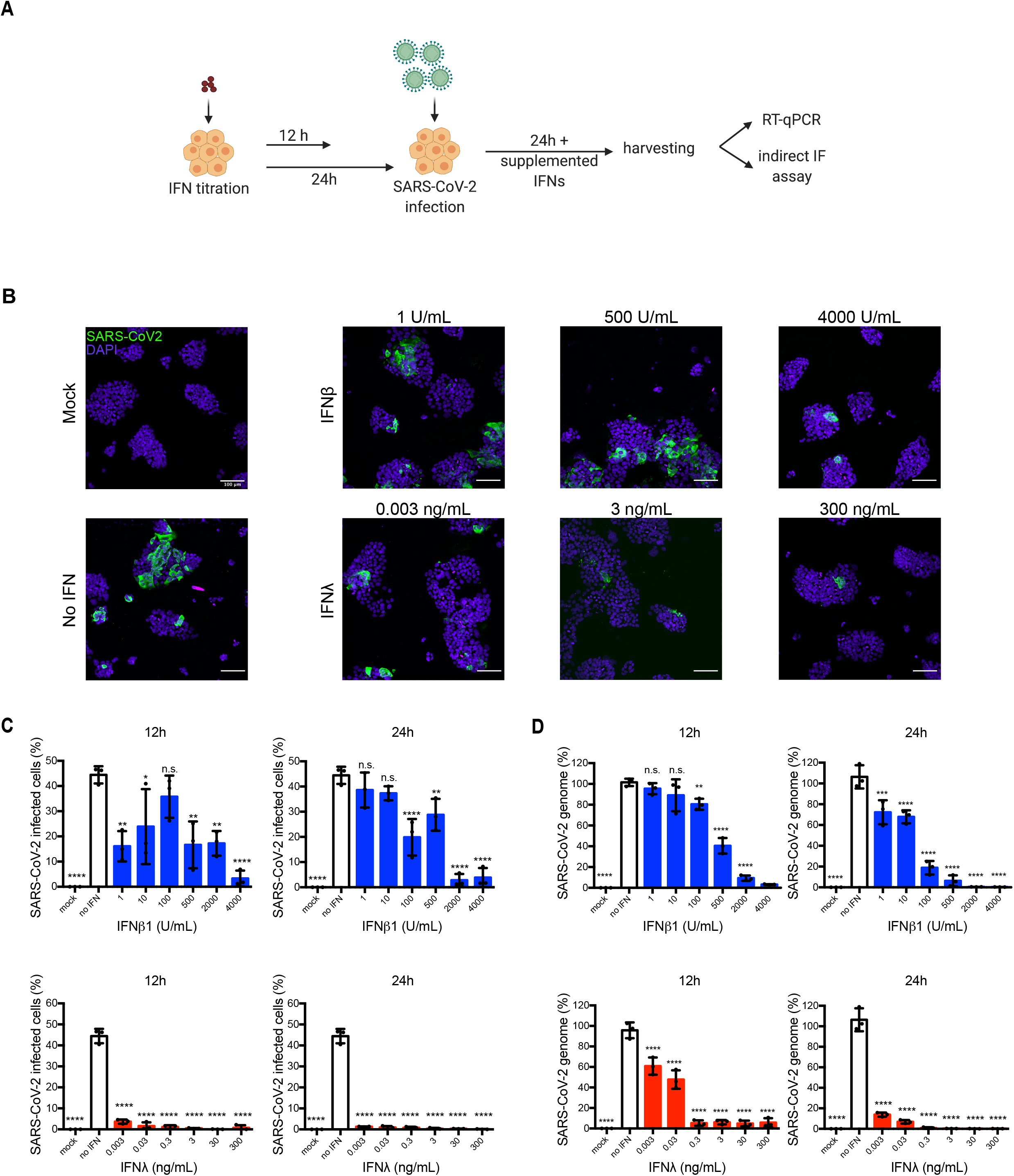
Exogenously added type I and III IFNs inhibit SARS-CoV-2 infection of hIECs in a concentration dependent manner. (A-D) WT T84 cells were mock-treated or pretreated with increasing concentration of type I (IFNβ1) and type III (IFNλ1/2/3) IFNs for 12 h and 24 h prior to infection. Cells were infected with SARS-CoV-2 using a MOI of 0.04. 24 hpi, cells were harvested to assay virus infection and replication. (A) Schematic of infection conditions. (B) Cells were fixed and indirect immunofluorescence was performed against the viral nucleocapsid (green). Nuclei were stained with DAPI (blue). Representative images are shown. Scale bars=100 μm. (C) The percentage of SARS-CoV-2-positive cells was quantified from images in (B). (D) RNA was harvested, and q-RT-PCR was used to evaluate the replication of the SARS-CoV-2 genome. Data are normalized to input and expressed as percentage, setting no-IFN treated cells to 100%. (C-D) Error bars indicate standard deviation. n = 3 biological replicates. n.s =not significant. P<0.05 *, P<0.01 **, P<0.001 ***, P <0.0001 **** as determined by ordinary one-way ANOVA with Dunnett’s multiple comparison test using non-treated infected cells as reference.

Taking in account that the units of measurement for type I and III IFNs are not comparable since IFNβ1 is in expressed as antiviral activity (U/mL) while IFNλ1/2/3 as weight (ng/mL), we were further interested to determine if these low type III IFN concentrations were acting with the same efficiency on other viruses. Vesicular stomatitis virus (VSV) is often used as a gold standard in virology (reviewed in (Munis et al., 2020)), therefore we performed the same pretreatment experiment using different IFN concentrations as for SARS-CoV-2 (Figure 2A) but with VSV expressing luciferase (VSV-Luc). T84 wild type cells were pretreated with either IFN as before for 24 or 12 h and then were subsequently infected with VSV-Luc. 8 hpi a luciferase assay was performed to determine virus infection levels. Interestingly, 0.0003 ng/mL were not enough to significantly inhibit VSV-Luc infection (Supplementary Figure 1B), while low concentrations of type I IFNs were efficient against the virus even with only 12 h pretreatment (Supplementary Figure 1A). This data suggests that type I IFNs are more efficient against VSV-Luc than type III IFNs as compared to SARS-CoV-2, which outlines the sensitivity of this coronavirus to type III IFNs.

Interestingly, when comparing the effects on type I and III IFN pretreatments on both the number of SARS-CoV-2 infected cells and genome replication, we could observe that longer IFN pretreatment and higher IFN concentrations were required to significantly reduce SARS-CoV-2 genome copy number when compared to the concentrations and time of pretreatment required to reduce the number of infected cells (Figure 2C-D). Such differences were especially significant for lower concentrations of IFN-treated cells for both IFNβ1 and IFNλ1/2/3. In order to find out the underlying reason of such an observation, we treated T84 wild-type cells with either 1U/mL of IFNβ1or 0.003 ng/mL of IFNλ1/2/3 for 12 hours prior to SARS-CoV-2 infection and compared them to the non-treated virus infected cells (Figure 3A). Cells were harvested 24hpi and immunostained using an antibody against the SARS-CoV-2 nucleoprotein or against double stranded RNA (J2) to monitor nucleoprotein expression and viral genome replication, respectively (Figure 3B). Quantification reveals that IFN mock-treated cells showed a similar number of nucleocapsid-positive cells and J2-positive cells while the ratio of J2-positive cells was higher than nucleocapsid-positive cells in both IFNβ1 and IFNλ1/2/3 treated cells (Figure 3C). Interestingly, this decrease in the number of cells positive for both dsRNA (J2 staining) and the SARS-CoV-2 nucleocapsid was more pronounced for the FN-λ1/2/3 treated cells (Figure 3B-C).

**Figure 3:**
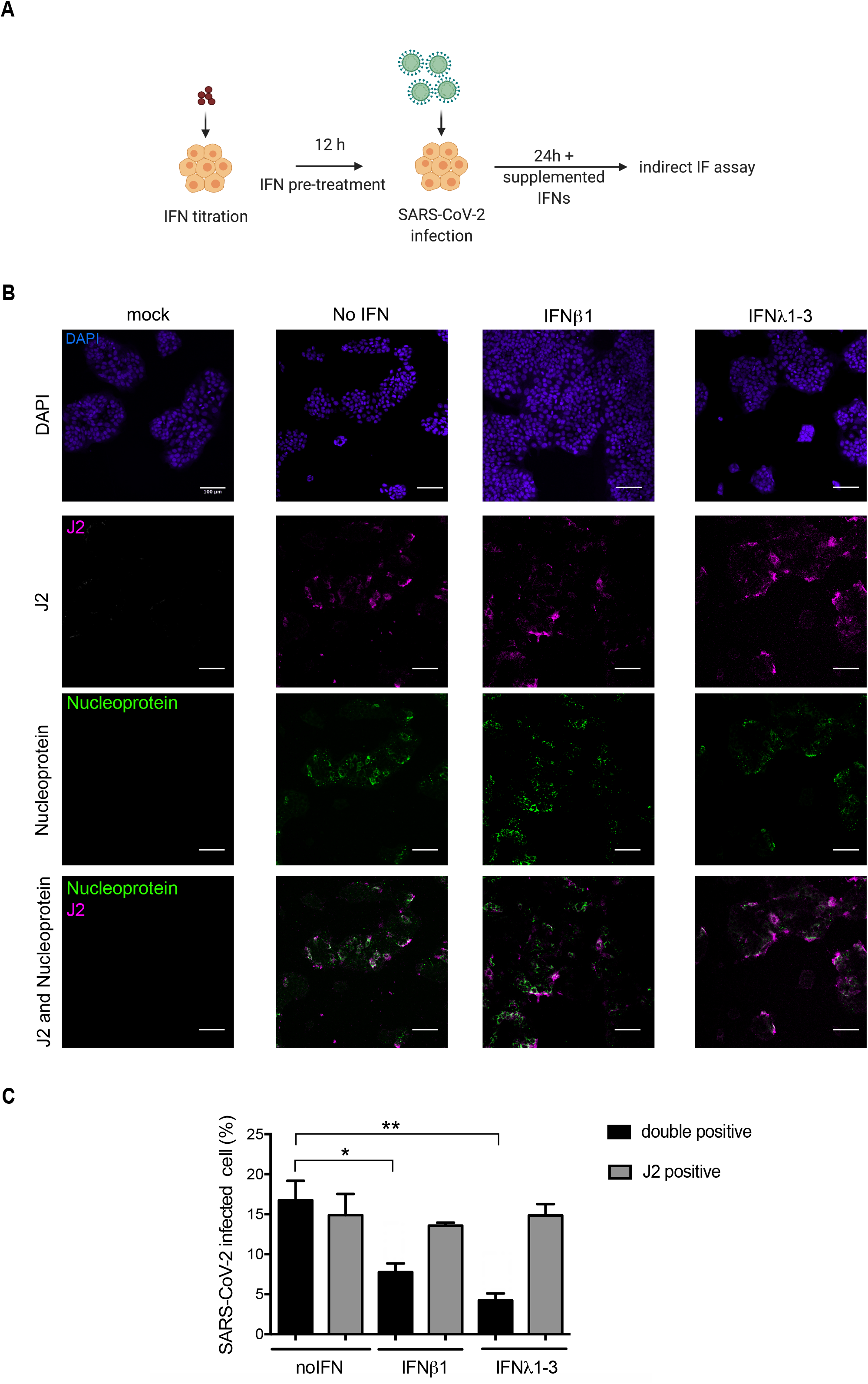
Type I and Type III IFNs induce an antiviral state to protect against SARS-CoV-2 in hIECs by interfering with nucleocapsid expression. (A-C) WT T84 cells were mock-treated or pretreated with type I and III IFN at low concentration (1 U/mL IFNβ1 or ng/mL IFNλ1/2/3) 12h prior to infection with SARS-CoV-2 using a MOI of 0.04. Cells were harvested 24 h post SARS-CoV-2 infection. (A) Schematic of infection set-up. (B) Cells were fixed, indirect immunofluorescence was performed against the viral nucleocapsid protein (green) and double stranded RNA (J2) (magenta), nuclei were stained with DAPI (blue). (C) The percentage of both nucleocapsid-positive and J2-positive cells (double-positive cells) and J2-positive only cells were quantified from (B). Error bars indicate standard deviation. n = 3 biological replicates. n.s= not significant. P<0.5 *, P<0.01 ** as determined by ordinary one-way ANOVA with Dunnett’s multiple comparison test to the non-treated infected cells as reference.

These results suggest that both type I and III IFNs are effective in mounting an antiviral immune response against SARS-CoV-2 not only by stopping virus genome replication but also by stalling or inhibiting nucleocapsid expression or translation. Differences between type I and type III IFNs suggest that the type III IFN is more effective in preventive SARS-CoV-2 protein translation compared to type I IFN.

### Pre-treatment with type III IFNs mediates a faster antiviral response against SARS-CoV-2 than pre-treatment with type I IFNs

To determine whether both type I and III IFNs require the same time to achieve a similar antiviral protection against SARS-CoV-2 in hIECs, we performed a time course experiment to determine the time required for each IFN to induce an antiviral state. Wild-type T84 cells were pretreated with either type I or type III IFN at different timepoints ranging from 24 hours to 3 hours prior to infection with SARS-CoV-2. Interferons were maintained throughout the time course of virus infection and viral genome load and the number of virus infected cells was determined by q-RT-PCR and immunofluorescence assay, respectively (Figure 4A). Initially, 4000 U/mL of IFNβ1 or 3ng/mL of IFNλ1/2/3 were used as 24h pre-treatment with these concentrations led to a complete inhibition of virus infection (Figure 2). Results show that both type I and type III IFNs display similar kinetics of antiviral activity when using high concentrations of either IFN (Figure 4B). Treating cells with either IFN for 3 hours was sufficient to completely deplete SARS-CoV-2 infection as determined by nucleocapsid immunostaining (Figure 4B, left panels). Viral genome was also greatly reduced with 3 hour pretreatment of either IFN, however only by around 60% as compared to no IFN-treated cells (Figure 4B, right panels). Moreover, even at long IFN-pretreatment times, for which no infected cells were determined by SARS-CoV-2 nucleocapsid immunostaining, viral genome could still be detected (Figure 4B). The detection limit of the antibody could lead to this phenotype, however this further supports our previous results suggesting that IFNs act by inhibiting virus translation (Figure 3).

**Figure 4:**
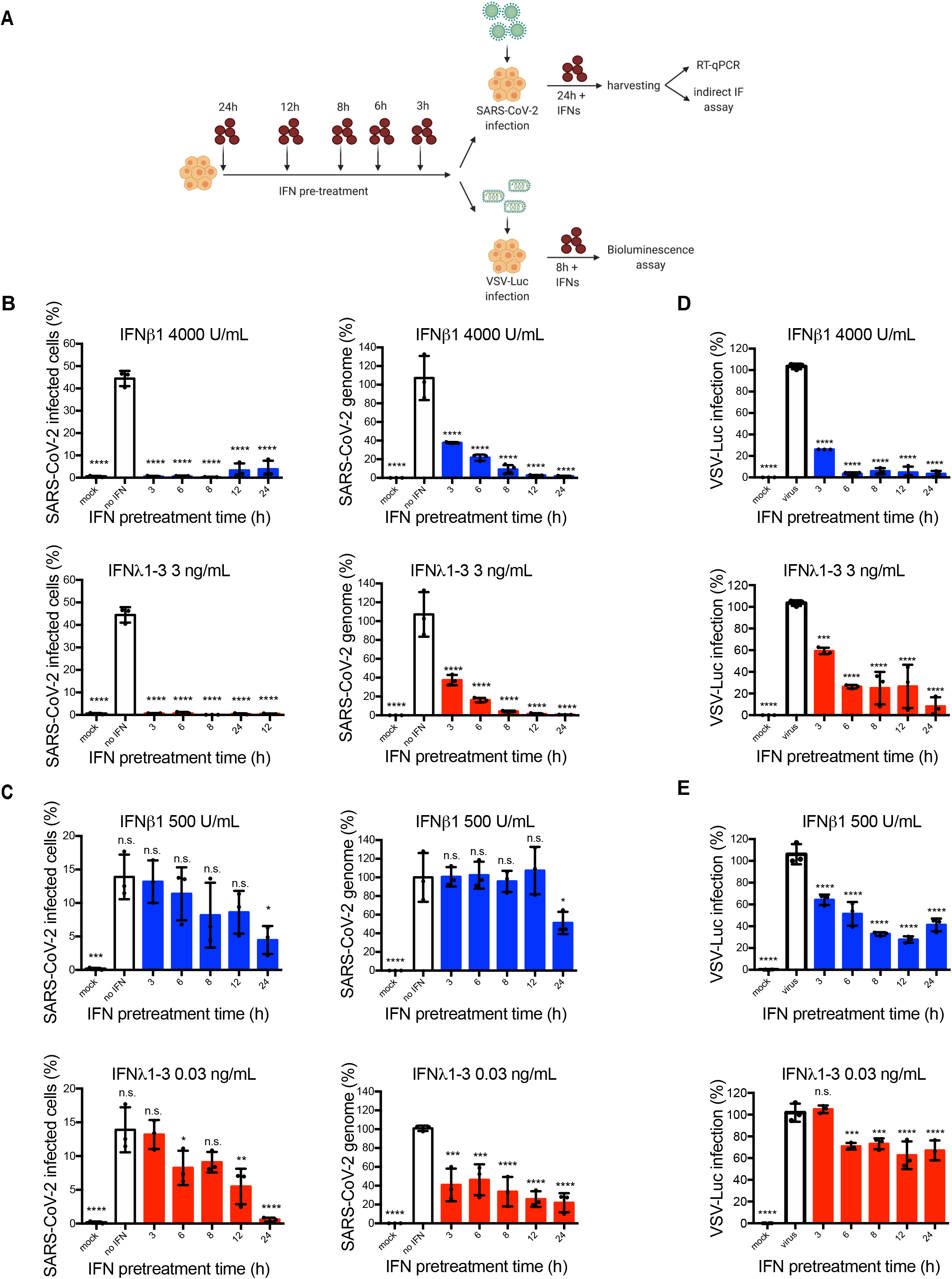
Differences between type I and type III IFNs in providing antiviral protection in hIECs against SARS-CoV-2 and VSV. (A-E) WT T84 cells were mock-treated or pretreated with type I and III IFN at high (4000 U/ml IFNβ1 or 3 IFNλ1/2/3) or low (500 U/ml IFNβ1 or 0.03 IFNλ1/2/3) concentrations for different timepoints prior to infection. Cells were infected with either SARS-CoV-2 using a MOI of 0.04 (as determined in Vero cells) or vesicular stomatitis virus expression Firefly luciferase (VSV-Luc) using a MOI of 5 (as determined in wild type T84 cells). 24 hpi (SARS-CoV-2) or 8 hpi (VSV-Luc) cells were harvested to assay virus infection. (A) Schematic of infection set-up. (B,C) The percentage of SARS-CoV-2-positive cells was quantified by immunofluorescence (left panels) and viral replication was addressed by q-RT-PCR (right panels). Q-RT-PCR data are normalized to input and expressed as percentage, setting no-IFN treated cells to 100%. (D,E) VSV-Luc infection was assayed by measuring the luciferase activity at 8 hpi. Luciferase activities for IFN treated samples were normalized to the Luciferase activity of the mock-treated sample, which correspond to 100%. (B-E) Error bars indicate standard deviation. n = 3 biological replicates. n.s.=not significant. P<0.5 *, P<0.01 **, P<0.001 ***, P <0.0001 **** as determined by ordinary one-way ANOVA with Dunnett’s multiple comparison test using non-treated infected cells as reference.

As the high IFN-concentrations were able to completely eliminate virus infection, we next addressed whether lower concentration of type I or type III IFNs would impact the kinetics of the antiviral program against SARS-CoV-2. T84 WT cells were pretreated with 500 U/mL of IFNβ1 or 0.03 ng/mL of IFNλ1/2/3 at different timepoints prior to infection with SARS-CoV-2 (Figure 4A). Results show that 24 hours pretreatment with low-concentration of type I IFN is required to significantly reduce both the number of SARS-CoV-2 infected cells and viral genome copies (Figure 4C). Shorter incubation times with type I IFNs does not lead to any decrease in the number of infected cells or the amount of virus genome copies (Figure 4C). On the contrary, shorter treatment times with type III IFNs lead to a decrease in both the number of infected cells and the amount of virus genome copies (Figure 4C). Furthermore, 24h pre-treatment with 0.03ng/ml of IFN-λ1/2/3 completely depletes the number of SARS-CoV-2 cells which are positive for nucleocapsid protein and significantly reduces the number of virus genome copies (Figure 4C). These results strongly suggest that type III IFN requires less time to establish an antiviral effect against SARS-CoV-2 compared to type I IFNs. As a note, we again observe that the viral genome is only reduced to 20% as compared to no IFN-treated cells when pretreating cells for 24 hour with type III IFNs, while viral nucleocapsid protein appears to be absent (Figure 4D, lower panels), suggesting a high detection limit of the antibody or an IFN-dependent inhibition of translation.

This faster kinetics of antiviral protection of type III IFNs is interesting as previous work has shown that type III IFNs require a longer time to establish their antiviral state (Bhushal et al., 2017; Kohli et al., 2012; Marcello et al., 2006; Meager et al., 2005; Pervolaraki et al., 2018; Sheppard et al., 2003). To determine if the fast type III IFN-induced antiviral effect against SARS-CoV-2 infection is virus specific, we used both high and low IFN concentrations and conducted the same time course experiment on T84 wild cells infected with VSV-Luc (Figure 4A). Similar to previous work, 8h pre-treatment of type I IFN at high concentrations was sufficient to reduce VSV infection by 90%, while type III IFN required 24h to achieve 90% reduction of infectivity (Figure 4D). Additionally, experiments performed using low IFN concentrations further supported that type I IFNs can control VSV infection faster than type III IFNs (Figure 4E). As such, 24 hours pre-treatment with type III IFNs reduced VSV-Luc infection only by 30% when compared to non-treated infected cells (Figure 4E). These results confirm that low concentrations of type III IFN require more time to establish an antiviral state to inhibit VSV-Luc infection compared to type I IFNs. Finally, when comparing the antiviral activities of low concentrations for both type I and type III IFNs against SARS-CoV-2 and VSV (Figure 4C *vs*. 4E), we could observe that SARS-CoV-2 is more resistant to type I IFN compared to VSV and, most importantly, more sensitive to type III IFNs compared to VSV.

Altogether these results suggest that type I IFNs require more time to establish an antiviral state in hIECs to control SARS-CoV-2 infection than type III IFNs. Interestingly, the potency and time-dependency of the IFN-induced antiviral state appears to also be virus specific with SARS-CoV-2 being particularly sensitive to type III IFNs.

### IFN pre- and post-treatment offers protection against SARS-CoV-2 spread

We have established that type III IFNs induce a better antiviral protection against SARS-CoV-2 compared to type I IFN when cells are treated prior to infection. To address whether type III IFNs can exercise their antiviral activities after viral infection, T84 WT cell were infected with SARS-CoV-2 at a MOI of 0.04 (as determined in Vero cells) and treated at different time points post-infection with either 4000 U/mL of IFNβ1 or 3 ng/mL of IFNλ1/2/3 (Figure 5A). 24 hpi virus genome copy number was evaluated and results show that both type I and III IFNs were able to inhibit viral replication when added up to 4h post-infection, and for type I IFN the effect was visible even when adding up 8h post-infection, demonstrating the effectiveness of IFN treatments against SARS-CoV-2 (Figure 5B).

**Figure 5:**
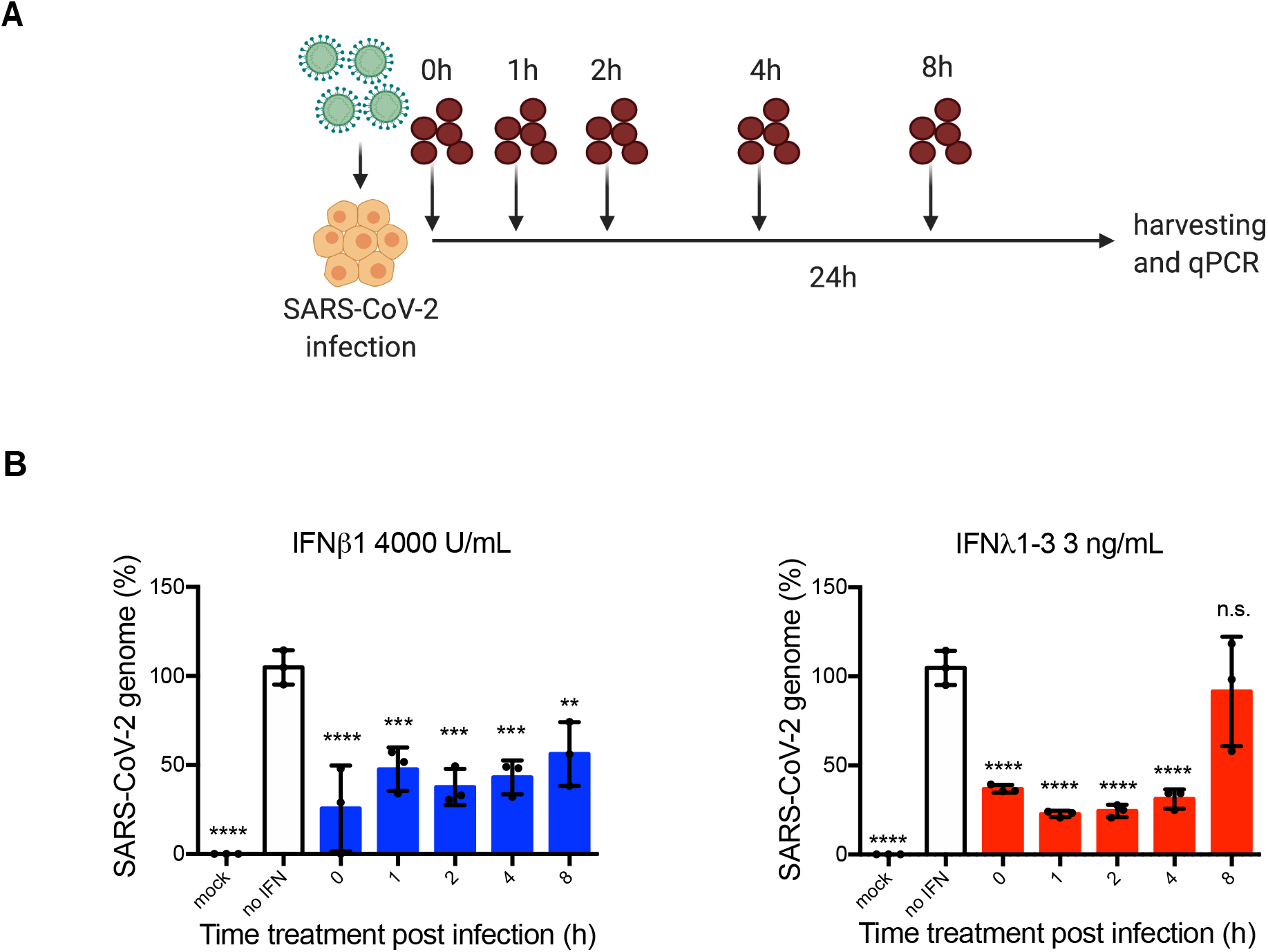
Kinetics of type I and III IFNs establishment of an antiviral state against SARS-CoV-2 in hIECs. (A-B) WT T84 cells were infected with SARS-CoV-2 using a MOI of (as determined in Vero cells). At 0, 1, 2, 4 or 8 hpi, 4000 U/mL of IFNβ1 or 3 ng/mL of IFNλ1/2/3 was added. (A) Schematic of experiment set-up. (B) 24 hpi, RNA was harvested to assess virus replication levels using q-RT-PCR, data are normalized to input and expressed as percentage, setting no-IFN treated cells to 100%. Error bars indicate standard deviation. n = 3 biological replicates. n.s.=not significant. P<0.5 *, P<0.01 **, P<0.001 ***, P <0.0001 **** as determined by ordinary one-way ANOVA with Dunnett’s multiple comparison test using non-treated infected cells as reference.

To determine how long the IFN-mediated antiviral state persists, T84 WT cells were treated with either 4000 U/mL of IFNβ1 or 3 ng/mL of IFNλ1/2/3 for 24 h, then cells were washed and fresh medium lacking IFNs was added. At 12, 24, 48 or 72 hours after medium exchange, WT T84 cells were infected with SARS-CoV-2 at a MOI of 0.04 (as determined in Vero cells) (Figure 6A). At 8 hpi, virus genome copy number was determined via q-RT-PCR and results showed that the antiviral state induced by both type I and III IFNs persisted up to 72 hours (Figure 6B). Interestingly, protection induced by type I IFN strongly decreased over time, 12h following media exchange of IFNβ1 treated cells, infection levels were reduced by 90%, 48h post-media exchange infection was reduced to 50%, and at 72h the antiviral effect was almost lost (Figure 6B). On the contrary, type III IFNs induce a longer lasting and more potent antiviral state. Even at 48h post-media exchange, virus replication was decreased by 75% and the antiviral effect was still present after 72h reducing the virus burden by 50% as compared to non-IFN-treated cells (Figure 6B). To see if this long term antiviral effect was SARS-CoV-2 specific, T84 wild type cells were infected with VSV-Luc following IFN washout (Figure 6A). Interestingly, the antiviral state which both type I and III IFNs induced to restrict this virus was less pronounced as compared to SARS-CoV-2 (Figure 6C). Importantly, the protection against VSV-Luc induced by type I IFNs was longer lasting than for type III IFNs, and reduced VSV-Luc infection significantly as compared to no IFN treatment even 24h post-media change (Figure 6C). On the contrary, for no timepoint post-media exchange the type III IFN-induced antiviral state was potent enough to significantly reduce VSV-Luc infection in hIECs (Figure 6C), thereby supporting the fact that the SARS-CoV-2 sensitivity to type III IFNs is virus specific.

**Figure 6:**
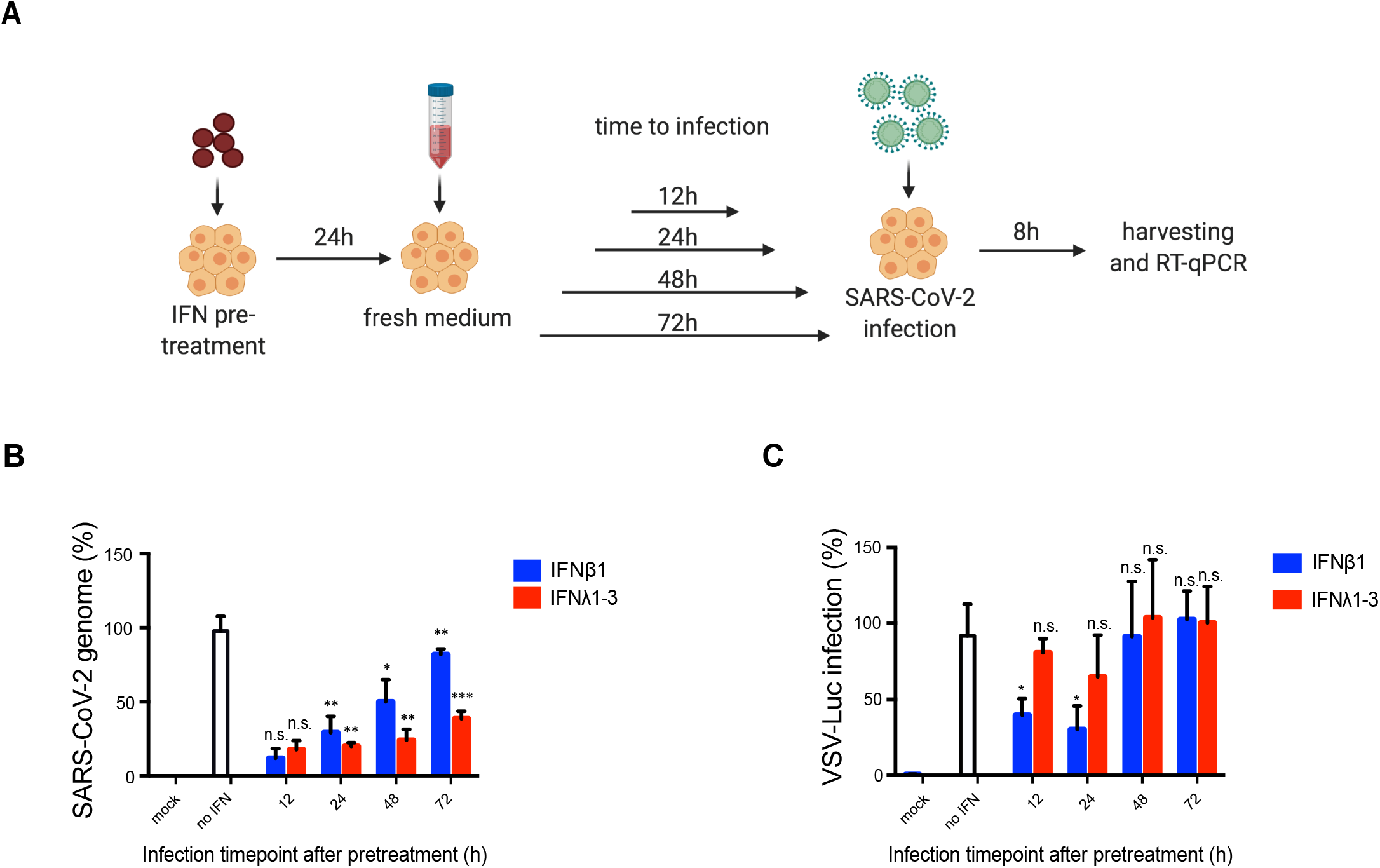
Type III IFNs induce a longer lasting antiviral state inhIEC against SARS-CoV-2 infection compared to type I IFN. (A-C) WT T84 cells were mock-treated or pretreated with 4000 U/mL of IFNβ1 or 3 ng/mL of IFNλ1/2/3. 24 hours after treatment, fresh medium was added to the cells. 12, 24, 48 or 72 hours after medium exchange, cells were infected. (A) Schematic of experiment set-up for SARS-CoV-2 infection. (B) Infection with SARS-CoV-2 using a MOI of 0.04 (as determined in Vero cells). 8 hpi, RNA was harvested to assess virus replication levels using q-RT-PCR, virus genome was normalized to input. (C) Infection with VSV-Luc using an MOI of 5 (as determined in T84 wildtype). 8 hpi, a Luciferase assay was performed to determine virus infection levels. (B-C) SARS-CoV-2 genome copy number or and VSV-Luc infection levels were normalized to the non-treated infected cells of the respective timepoint. Error bars indicate standard deviation. n = 3 biological replicates. n.s.=not significant. P<0.5 *, P<0.01 **, P<0.001 ***, P <0.0001 **** as determine by a two-tailed unpaired t-test with Welch’s correlation, using non-treated infected cells of the respective time point as reference.

Taken together, both IFNs are fast acting and can prevent viral replication and spread even when added after SARS-CoV-2 infection took place. Furthermore, our data suggests that type I and III IFNs can induce an antiviral state that lasts for several days. Type III IFNs are especially potent and long lasting as they can inhibit SARS-CoV-2 more efficiently compared to type I IFN.

## Discussion

There is increasing evidence that SARS-CoV-2 infection is not restricted to the respiratory tract but also impacts other organs (Trypsteen et al., 2020) as viral components such as RNA and proteins were detected in patient biopsies or in postmortem tissues such as the heart (Remmelink et al., 2020; Tavazzi et al., 2020), kidney (Bradley et al., 2020), brain (Bradley et al., 2020; Puelles et al., 2020; Schaller et al., 2020) and more. The gastrointestinal tract is one such important secondary organs and a great fraction of COVID-19 patients display gastrointestinal symptoms and shed viral genomes within their feces (Guan et al., 2020; Huang et al., 2020; L. Lin et al., 2020; Wölfel et al., 2020; Y. Wu et al., 2020; Xiao et al., 2020; Xing et al., 2020; Xu et al., 2020). This highlights the importance of understanding the molecular interaction of this specific virus with the gastrointestinal tract in order to better characterize and treat the associated pathogenesis and to curb the pandemic efficiently. Together our results have shown that both type I and type III IFNs are efficient in mounting an antiviral immune response against SARS-CoV-2 in human intestinal epithelial cells as seen by the rapid virus replication and spread in the absence of either type I or type III IFN receptors (Figure 1). These observations highlight the significance of type I and type III IFN signaling during SARS-CoV-2 infection in the human intestinal epithelium as opposed to murine models of SARS-CoV-2 infection, where type I IFN only minimally restricts SARS-CoV-2 (Israelow et al., 2020). The importance of both IFNs in controlling SARS-CoV-2 in human tissue has been supported by several other studies (Lamers et al., 2020; Stanifer, Kee, et al., 2020; Triana et al., 2021; Zang et al., 2020), however the differences between the two types of IFNs has yet to be explored. It is important to point out that our results showed differences between the two types of IFNs in many ways. Firstly, we describe an interesting observation where type I IFN appears to be more important in controlling early steps of the SARS-CoV-2 infection cycle since T84 IFNAR^−/−^ cells were more infected at early timepoints when compared to T84 wildtype cells, while type III IFNs play a more critical role in stopping viral replication and spread in human intestinal epithelium as seen from high infection and viral genome rate in T84 IFNLR^−/−^ cells at late timepoints (Figure 1).

These results point out that there are indeed differences in the kinetics of antiviral properties of type I and III interferons against SARS-CoV-2 infections in human intestinal epithelial cells. This however, remains unclear in human lung epithelium as there are contradictory observations if there is production of type I and III IFNs in the cells during SARS-CoV-2 infection (Blanco-Melo et al., 2020; Chu et al., 2021; Desai et al., 2020; Katsura et al., 2020; Shuai et al., 2020; Vanderheiden et al., 2020). To the best of our knowledge, type III IFNs have been vastly neglected, although some studies did indeed show that the human lung epithelium is also sensitive to type III IFN treatment (Busnadiego et al., 2020; Felgenhauer et al., 2020; Vanderheiden et al., 2020). Importantly, no similar direct comparison of the antiviral activities of both type I and type III IFNs has been performed in human lung epithelial cells. Interestingly, inhibition of *de novo* infectious virus production in Calu-3 cells suggests that type I IFN would be more potent as compared to type III IFNs (Felgenhauer et al., 2020). This further highlights the importance of studying SARS-CoV-2 in multiple cellular models covering the broad tropism of SARS-CoV-2 as the sensitivity of this pathogen to various antiviral strategies might be organ specific.

We showed by immunostaining of the virus nucleocapsid protein and quantification of the viral genome that both type I and III IFNs restrict SARS-CoV-2 in a concentration-dependent manner. Interestingly, inhibition of nucleocapsid protein with a certain IFN concentration did not mirror the inhibition of the virus genome, since the genome load was still measured in samples where no nucleocapsid could be detected. To analyze in more detail the discrepancy between the viral genome copy number and number of infected we further determined whether infected and treated cells were positive for J2, which labels dsRNA and is representative for virus transcription, and nucleocapsid protein representing virus translation. We have shown that in contrast to SARS-CoV-2 infected cells without treatment, having similar levels of viral double stranded RNA and nucleocapsid protein, IFN-treated cells can be positive for viral double stranded RNA while no nucleocapsid protein could be detected. This observation is especially prominent in type III IFN-treated cells and strongly suggests that the underlying mechanism of IFNs to restrict virus spread is by inhibiting virus translation. Previously it was already reported that type I IFN-treatment during Dengue and HIV infection can affect virus protein translation (Coccia et al., 1994; Diamond & Harris, 2001). However, little is known about type I and especially type III IFN-mediated inhibition of coronavirus nucleocapsid translation, and specifically for SARS-CoV-2-infection. Our observation suggests a possible mechanism on the mode of action of IFN-dependent signaling on SARS-CoV-2 and shows that type I and III IFNs efficiently restrict the spread of SARS-CoV-2 in human epithelium after virus entry by inhibiting the production of nucleocapsid (and probably other viral proteins) and thus reducing the release of functional virus particles to neighboring cells. More needs to be done in order to decipher the details of this mechanism underlying the IFN-mediated inhibition of viral genome translation. Indeed, several interferon stimulated genes (ISGs) were found to inhibit viral protein translation and might be good candidates as key cellular players in the IFN-dependent response to SARS-CoV-2. A well investigated type I and III-induced ISG is PKR, which upon activation phosphorylates EIF2α to halt cellular translation (Sadler & Williams, 2008). Also the IFIT family was demonstrated to suppress cellular translation upon virus infections by several mechanisms (Diamond & Farzan, 2013; Vladimer et al., 2014). Moreover, ISG20 was shown to impair translation of virus protein without affecting cellular protein translation by discriminating self from non-self (N. Wu et al., 2019).

Our data supports that type I and type III IFNs play different roles at different stages of virus infection due to their differences in their kinetics of antiviral action. While both IFN treatments induce a dose and time dependent inhibition of viral infection in hIECs, our results strongly suggest that type III IFNs were able to restrict virus infection at lower dosage and with shorter treatment time as compared to type I IFNs. Comparing whether type I or type III IFNs are more potent antiviral at same the concentrations is intrinsically difficult as the IFNs are available at concentration expressed in different units. The type III IFNs are available in protein concentration (ng/mL) while the type I IFN concentration is expressed as an antiviral activity (U/mL). Interestingly, our observations suggest that type III IFNs are more potent and faster acting on SARS-COV-2 is contradictory to a previous study using VSV infection of hIECs (Pervolaraki et al., 2018), which shows that type III IFNs need longer time to confer protection. To address these discrepancies, we here compared the effect of both type I and type III IFNs on VSV and SARS-CoV-2 infection (Figure 2, Supplementary Figure 1, Figure 4). We could reproduce our previous results that type I IFN was more potent in inhibiting VSV infection especially at lower concentrations (Supplementary Figure 1A, Figure 4D-E). Interestingly, by comparing SARS-CoV-2 and VSV infections, we could show that low concentration of type I IFN was able to control VSV infection while having little to no effect on SARS-CoV-2 infection (Figure 2B-C, Supplementary Figure 1A, Figure 4). Reciprocally, low concentration of type III IFNs were able to control SARS-CoV-2 but with limited to no impact on VSV (Figure 1, Supplementary Figure 1B, Figure 4). These differences in the efficacy of type I and type III IFNs in providing, in the same cell type, an antiviral protection against two distinct viruses suggest that two distinct antiviral states are achieved upon type I and type III IFN treatment and that these states are more potent against specific viruses. While there is very limited evidence that type I and type III IFN induce the expression of different ISGs (Selvakumar et al., 2017), it recently became clear that both cytokines induce the same ISG but with very different kinetics as such, likely creating distinct antiviral state (Crotta et al., 2013; Jilg et al., 2014; J. Da Lin et al., 2016; Marcello et al., 2006; Pervolaraki et al., 2018; Voigt & Yin, 2015; Zhou et al., 2007).

Lastly, we have demonstrated that addition of IFNs after SARS-CoV-2 infection was still able to inhibit viral replication (Figure 5) which further highlights the effectiveness of IFN treatments against SARS-CoV-2 infection. Importantly, we could show that type III IFNs were able to provide longer-lasting protection against SARS-CoV-2 compared to type I IFNs and such potent antiviral states still persist and are able to inhibit virus replication for more than 72 hours upon withdrawal (Figure 6B). This effect is specific for SARS-CoV-2, since no such long-lasting effect was determined upon VSV-Luc infection (Figure 6C). Altogether we see that type III IFNs act fast, require low concentration and short pre-treatment time, and offer long-lasting antiviral protection against SARS-CoV-2 in hIECs. These observations suggest type III IFN treatment as a strong therapeutic candidate against SARS-CoV-2 infection of the human intestine.

It has been previously suggested to use IFNβ alone or in combination with other antiviral agents to treat SARS-CoV-1 infection (Cinatl et al., 2003). Furthermore, clinical trials using treatment combining intravenous injection of IFNβ1 and lopinavir/ritonavir were performed in Saudi Arabia to treat MERS-CoV (Arabi et al., 2018, 2020). IFNβ1 seems to be the most efficient interferon to curb coronavirus infection, and a report shows that it is due to the fact that IFNβ1 can induce the production of adenosine with anti-inflammatory properties and maintain endothelial barrier function in pulmonary endothelial cells via upregulation of cluster of differentiation 73 (CD73) (Sallard et al., 2020). SARS-CoV-2 is thought to be more sensitive to IFN treatment due to its truncated Orf6 and Orf3 protein as opposed to those of SARS-CoV-1 and MERS-CoV due to a loss of their inhibitory effects on IFNs signaling pathway (Felgenhauer et al., 2020; Lokugamage et al., 2020). There is currently a treatment consisting of a triple combination of IFNβ-1b, lopinavir-ritonavir and ribavirin to treat patients with COVID-19 in Hong Kong as a phase-2 trial (Hung et al., 2020). This trial has yielded promising outcomes showing symptoms alleviation and reduction of viral shedding duration for patients with mild to moderate COVID-19 disease in the hospital (Hung et al., 2020). Another study also pointed out that IFNβ-1a, when administered at clinically permissible concentration after SARS-CoV-2 infection, was highly effective in inhibiting *in vitro* SARS-CoV-2 replication (Clementi et al., 2020).

Here, we have demonstrated that, as compared to type I IFNs, type III IFNs are more potent in protecting human intestinal epithelial cells against SARS-CoV-2. This places type III IFNs as a promising option in treating SARS-CoV-2 infection particularly in the context of the human intestinal epithelium. Given the lower dose and shorter treatment required to confer an antiviral state, as well as the fact that the type III IFN antiviral activity persists longer than the type I IFN-mediated one, it is likely that the administration of type III IFN to patients would be more beneficial and would require a less frequent regimen which will render this approach more amenable as therapeutic option. Importantly and as reviewed before (Biggioggero et al., 2010), type I IFN treatment was shown to support development of autoimmune diseases and can induce tissue damage. We believe that type III IFNs could potentially be the superior choice of treatment than type I IFNs in treating SARS-CoV-2 in patients.

## Acknowledgements

This work was supported by research grants from the Deutsche Forschungsgemeinschaft (DFG): project numbers 415089553 (Heisenberg program), 240245660 (SFB1129), 278001972 (TRR186), and 272983813 (TRR179), the state of Baden Wuerttemberg (AZ: 33.7533.-6-21/5/1) and the Bundesministerium Bildung und Forschung (BMBF) (01KI20198A) to SB. MS was supported by the DFG (416072091) and the BMBF (01KI20239B). CMZ is supported by the SFB1129 (240245660). We also acknowledge funding from the Helmholtz International Graduate School for Cancer Research to CK, the German Academic Exchange Service (DAAD) (Research Grant 57440921) to PD, and the China Scholarship Council and the Landesgraduirtenförderung (LGF) to CG.

## Author Contributions

CMZ, CK, PD, performed experiments, analyzed data, and helped with manuscript writing. CG performed the VSV experiments. SB and MLS conceived experiments, interpreted results and wrote the manuscript. The final version of the manuscript was approved by all authors.

## Declaration of Interests

The authors declare no competing interests.

## Materials and Methods

### Cell line and Viruses

Wild type T84 (ATCC CCL-248) and their IFN receptor knock-outs (Pervolaraki et al., 2017) were cultured in a 50:50 mixture of Dulbecco’s modified Eagle’s medium (DMEM) and F12 (Gibco) supplemented with 10% fetal bovine serum and 1% penicillin/streptomycin (Gibco). T84 Wild-type and IFN receptor knock-out cell lines carrying the H2B fluorescent nuclear tag were generated by lentiviral transduction and antibiotic selection. Vero E6 cells (ATCC CRL 1586) were cultured in Dulbecco’s modified Eagle’s medium (DMEM) (Gibco) supplemented with 10% fetal bovine serum and 1% penicillin/streptomycin (Gibco).

SARS-CoV-2 was isolated from an infected patient at the University Hospital Heidelberg. The virus was amplified in Vero E6 cells and P3 virus stocks were used in all experiments (Stanifer, Kee, et al., 2020). VSV-Luc was prepared and used as previously described (Pervolaraki et al., 2018).

### Viral infections

All SARS-CoV-2 infections were performed with a multiplicity of infection of 0.04 as determined in Vero E6 cells. Prior to infection, culture media was removed and virus was added to cells and incubated for 1 hour at 37°C. Following the incubation, virus was removed and fresh media or media supplemented with indicated interferons was added back to the cells upon virus removal.

### Interferons treatment

Human recombinant IFN-beta 1a (IFNb) was obtained from Biomol (#86421). Recombinant human IFNλ 1 (IL-29) (#300-02L), IFNλ 2 (IL28A) (#300-2K) and IFNλ 3 (IL-28B) (#300-2K) were purchased from Peprotech. The IFN concentrations used to treat the cells and the duration of the treatments are stated in the figure legends.

### RNA isolation, cDNA synthesis and q-RT-PCR

Cells were harvested either 4, 8, 12 or 24 hours post infection and RNA was isolated using RNAeasy RNA extraction kit (Qiagen) as per manufacturer’s instructions. Complementary DNA was synthesized using iSCRIPT reverse transcriptase (BioRad) from 250 ng of total RNA per 20μL reaction according to the manufacturer’s instructions. Quantitative RT-PCR assay was performed using iTaq SYBR green (BioRad) as per manufacturer’s instructions. The expression of target genes was normalized to endogenous control *TBP.* Primer sequences are as below:

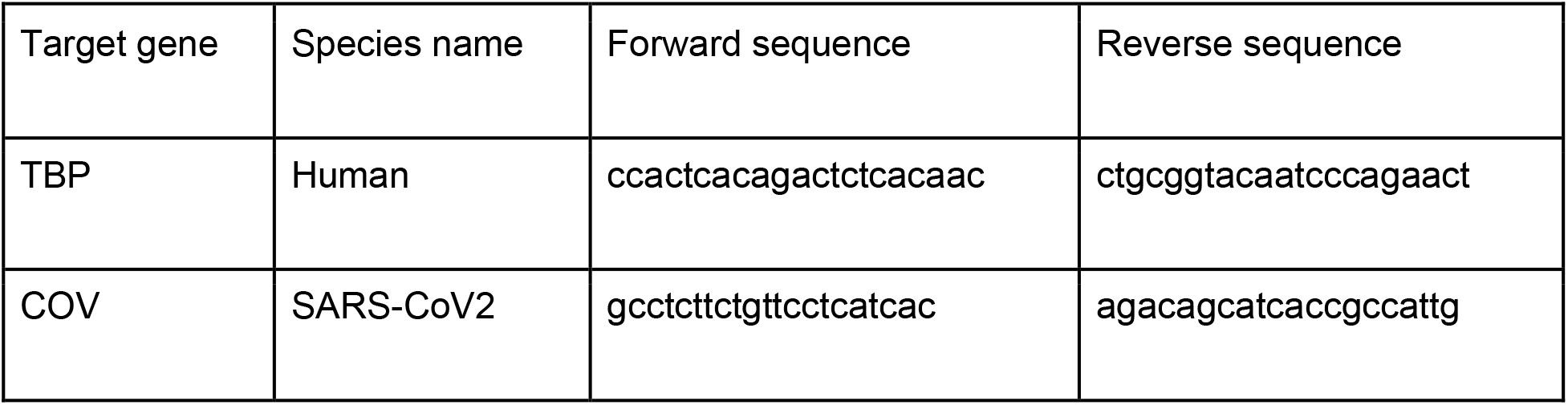

The fold change in SARS-CoV-2 genome copy number was calculated using input as a reference. Input RNA was harvested immediately after the 1-hour SARS-CoV-2 infection.

### Western blot

Cells were harvested and lysed with 1X RIPA (150 mM sodium chloride, 1.0% Triton X-100, 0.5% sodium deoxycholate, 0.1% sodium dodecyl sulphate (SDS), 50 mM Tris, pH 8.0) with phosphatase and protease inhibitors (Sigma-Aldrich) for 5 mins at 37°C. Lysates were collected and equal protein amounts were separated by SDS-PAGE and blotted onto a nitrocellulose membrane by wet-blotting (Bio-Rad). Membranes were blocked with TBS-T containing 5% Bovine Serum Albumin (BSA) for two hours at room temperature (RT). Primary antibodies beta-Actin (Sigma #5441) and phospho STAT1 (BD Transductions #612233) were diluted in the same blocking buffer and incubated overnight at 4°C. Membranes were then washed three times with TBS-T for 10 mins at RT with rocking. Anti-mouse antibodies coupled with horseradish peroxidase (GE Healthcare #NA934V) were used at 1:5000 dilution in blocking buffer and incubated at RT for 1 hour with rocking. Membranes were washed three times with TBS-T for 10 mins at RT with rocking. HRP detection reagent (GE Healthcare) was mixed 1:1 and incubated at RT for 5mins. Membranes were exposed to film and developed.

### VSV luciferase assay

Wild type T84 cells were seeded in a black F-bottom 96-well plate. Cells were treated as indicated in the text with type I or type III IFNs. VSV-luc was added to the wells using a MOI of 5 as determined in wild type T84 cells and the infection was allowed to proceed for 8 hours. At the end of the infection, media was removed, cells were washed 1X with PBS and lysed with Cell Lysis Buffer (Promega) at RT for 20 mins. A 1:1 dilution of Steady Glo (Promega) and Lysis Buffer were added to the cells and incubated at RT for 15 mins. Luminescence was read using an Omega Luminometer.

### Indirect Immunofluorescence Assay

Cells were seeded on iBIDI glass bottom 8-well chamber slides previously coated with 2.5% human collagen in water. At indicated times post-infection, cells were fixed in 4% paraformaldehyde (PFA) for 20 mins at RT. Cells were washed in 1X PBS and permeabilized in 0.5% Triton-X for 15 mins at RT. Cells were blocked using 3% BSA-PBS for 30 mins at RT. Mouse monoclonal antibody against SARS-CoV-2 Nucleocapsid protein (Sino biologicals MM05) was diluted in 1% BSA-phosphate-buffered saline (PBS) and incubated for 1h at RT. Cells were washed with 1X PBS three times and incubated with secondary antibodies conjugated with AF488 or AF647568 (Molecular Probes) and DAPI for 30-45 mins at RT. Cells were washed in 1X PBS three times and maintained in PBS. Cells were imaged on a Nikon/Andor Spinning Disc Confocal microscope to quantify the number of infected cells relative to the number of nuclei.

### Statistics and computational analyses and statistics

All statistical analysis was performed either by Ordinary one way ANOVA Dunnett’s multiple comparison test or by a two-tailed unpaired t-test with Welch’s correlation (specified in figure legend) using the GraphPad Prism software package.

In order to quantify infected cells from indirect immunofluorescent stained samples, ilastik 1.2.0 was used on DAPI images to generate a mask representing each nucleus as an individual object. These masks were used on CellProfiler 3.1.9 to measure the intensity of the conjugated secondary antibodies in each nucleus. A threshold was set based on the basal fluorescence of non infected samples, and all nuclei with a higher fluorescence were considered infected cells.

---

**Supplementary Figure 1:**
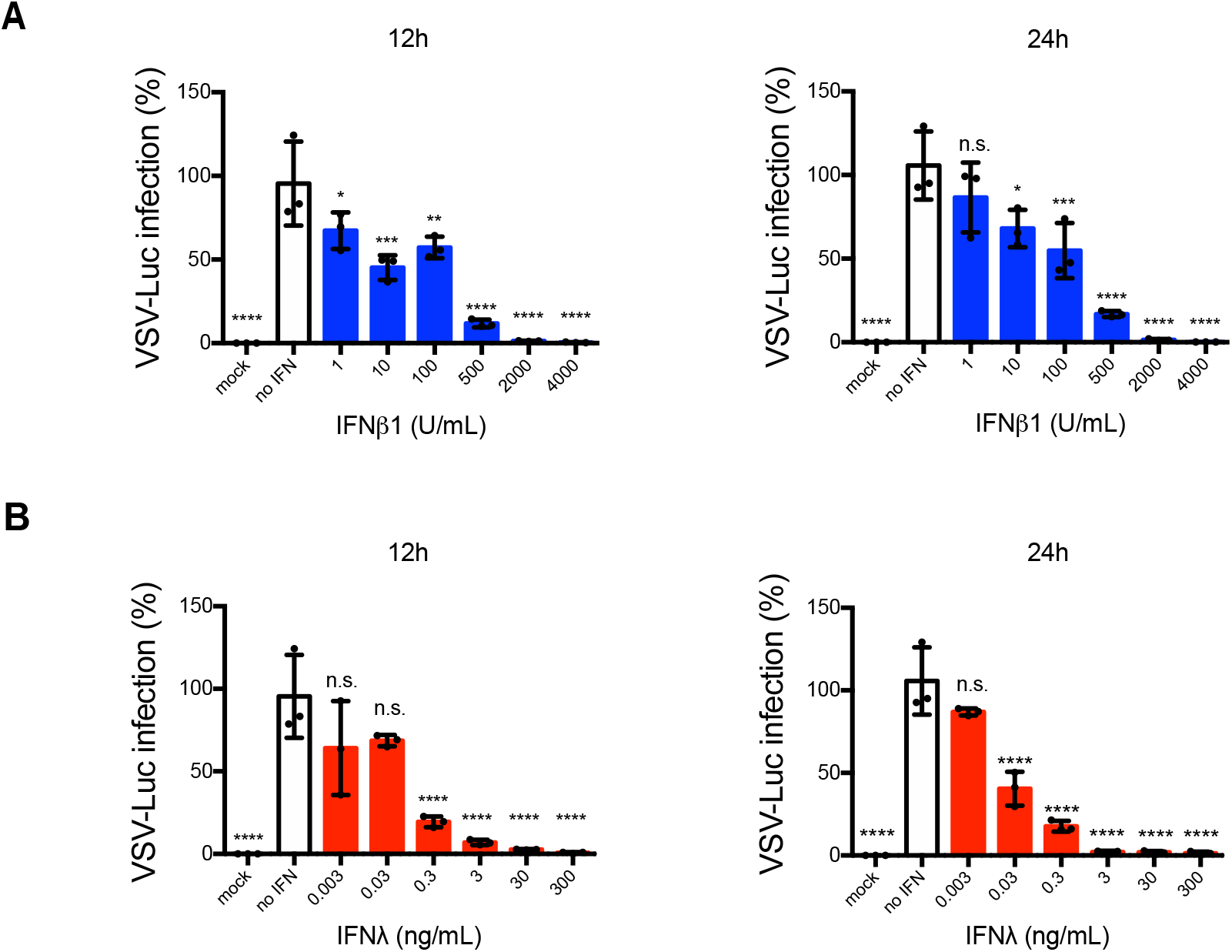
Exogenously added type I and III IFNs inhibit VSV-Luc infection of hIECs in a concentration dependent manner. (A-B) WT T84 cells were mock-treated or pretreated with increasing concentration of (A) type I (IFNβ1) and (B) type III (IFNλ1/2/3) IFNs for 12 h and 24 h prior to infection. Cells were infected with VSV-Luc using a MOI of 5 (as determined in T84 wildtype). 8 hpi, cells were harvested to assay virus infection with a Luciferase assay. The VSV-Luc infection was quantified and expressed as percentage, setting no-IFN treated cells to 100%. Error bars indicate standard deviation. n = 3 biological replicates. n.s.=not significant. P<0.05 *, P<0.01 **, P<0.001 ***, P <0.0001 **** as determined by ordinary one-way ANOVA with Dunnett’s multiple comparison test using non-treated infected cells as reference.

## References

Arabi, Y. M., Asiri, A. Y., Assiri, A. M., Balkhy, H. H., Al Bshabshe, A., Al Jeraisy, M., Mandourah, Y., Azzam, M. H. A., Bin Eshaq, A. M., Al Johani, S., Al Harbi, S., Jokhdar, H. A., Deeb, A. M., Memish, Z. A., Jose, J., Ghazal, S., Al Faraj, S., Al Mekhlafi, G. A., Sherbeeni, N. M., … Alothman, A. (2020). Interferon Beta-1b and Lopinavir–Ritonavir for Middle East Respiratory Syndrome. New England Journal of Medicine, 383(17), 1645–1656. https://doi.org/10.1056/nejmoa2015294

Arabi, Y. M., Mandourah, Y., Al-Hameed, F., Sindi, A. A., Almekhlafi, G. A., Hussein, M. A., Jose, J., Pinto, R., Al-Omari, A., Kharaba, A., Almotairi, A., Al Khatib, K., Alraddadi, B., Shalhoub, S., Abdulmomen, A., Qushmaq, I., Mady, A., Mady, O., Al-Aithan, A. M., … Fowler, R. A. (2018). Corticosteroid therapy for critically ill patients with middle east respiratory syndrome. American Journal of Respiratory and Critical Care Medicine, 197(6), 757–767. https://doi.org/10.1164/rccm.201706-1172OC

Baldridge, M. T., Nice, T. J., McCune, B. T., Yokoyama, C. C., Kambal, A., Wheadon, M., Diamond, M. S., Ivanova, Y., Artyomov, M., & Virgin, H. W. (2015). Commensal microbes and interferon-λ determine persistence of enteric murine norovirus infection. Science, 347(6219), 266–269. https://doi.org/10.1126/science.1258025

Bhushal, S., Wolfsmüller, M., Selvakumar, T. A., Kemper, L., Wirth, D., Hornef, M. W., Hauser, H., & Köster, M. (2017). Cell polarization and epigenetic status shape the heterogeneous response to type III interferons in intestinal epithelial cells. Frontiers in Immunology, 8(JUN). https://doi.org/10.3389/fimmu.2017.00671

Biggioggero, M., Gabbriellini, L., & Meroni, P. L. (2010). Type I interferon therapy and its role in autoimmunity. Autoimmunity, 43(3), 248–254. https://doi.org/10.3109/08916930903510971

Blanco-Melo, D., Nilsson-Payant, B. E., Liu, W. C., Uhl, S., Hoagland, D., Møller, R., Jordan, T. X., Oishi, K., Panis, M., Sachs, D., Wang, T. T., Schwartz, R. E., Lim, J. K., Albrecht, R. A., & tenOever, B. R. (2020). Imbalanced Host Response to SARS-CoV-2 Drives Development of COVID-19. Cell, 181(5), 1036–1045.e9. https://doi.org/10.1016/j.cell.2020.04.026

Bradley, B. T., Maioli, H., Johnston, R., Chaudhry, I., Fink, S. L., Xu, H., Najafian, B., Deutsch, G., Lacy, J. M., Williams, T., Yarid, N., & Marshall, D. A. (2020). Histopathology and ultrastructural findings of fatal COVID-19 infections in Washington State: a case series. The Lancet, 396(10247), 320–332. https://doi.org/10.1016/S0140-6736(20)31305-2

Busnadiego, I., Fernbach, S., Pohl, M. O., Karakus, U., Huber, M., Trkola, A., Stertz, S., & Hale, A. G. (2020). Antiviral activity of type i, ii, and iii interferons counterbalances ace2 inducibility and restricts sars-cov-2. MBio, 11(5), 1–10. https://doi.org/10.1128/mBio.01928-20

Chu, H., Chan, J. F. W., Wang, Y., Yuen, T. T. T., Chai, Y., Shuai, H., Yang, D., Hu, B., Huang, X., Zhang, X., Hou, Y., Cai, J. P., Zhang, A. J., Zhou, J., Yuan, S., To, K. K. W., Hung, I. F. N., Cheung, T. T., Ng, A. T. L., … Yuen, K. Y. (2021). SARS-CoV-2 Induces a More Robust Innate Immune Response and Replicates Less Efficiently Than SARS-CoV in the Human Intestines: An Ex Vivo Study With Implications on Pathogenesis of COVID-19. CMGH, 11(3), 771–781. https://doi.org/10.1016/j.jcmgh.2020.09.017

Cinatl, J., Morgenstern, B., Bauer, G., Chandra, P., Rabenau, H., & Doerr, H. W. (2003). Treatment of SARS with human interferons. Lancet, 362(9380), 293–294. https://doi.org/10.1016/S0140-6736(03)13973-6

Clementi, N., Ferrarese, R., Criscuolo, E., Diotti, R. A., Castelli, M., Scagnolari, C., Burioni, R., Antonelli, G., Clementi, M., & Mancini, N. (2020). Interferon-β-1a inhibition of severe acute respiratory syndrome-coronavirus 2 in vitro when administered after virus infection. Journal of Infectious Diseases, 222(5), 722–725. https://doi.org/10.1093/infdis/jiaa350

Coccia, E. M., Krust, B., & Hovanessian, A. G. (1994). Specific inhibition of viral protein synthesis in HIV-infected cells in response to interferon treatment. Journal of Biological Chemistry, 269(37), 23087–23094. https://doi.org/10.1016/s0021-9258(17)31623-x

Crotta, S., Davidson, S., Mahlakoiv, T., Desmet, C. J., Buckwalter, M. R., Albert, M. L., Staeheli, P., & Wack, A. (2013). Type I and Type III Interferons Drive Redundant Amplification Loops to Induce a Transcriptional Signature in Influenza-Infected Airway Epithelia. PLoS Pathogens, 9(11). https://doi.org/10.1371/journal.ppat.1003773

Desai, N., Neyaz, A., Szabolcs, A., Shih, A. R., Chen, J. H., Thapar, V., Nieman, L. T., Solovyov, A., Mehta, A., Lieb, D. J., Kulkarni, A. S., Jaicks, C., Xu, K. H., Raabe, M. J., Pinto, C. J., Juric, D., Chebib, I., Colvin, R. B., Kim, A. Y., … Deshpande, V. (2020). Temporal and spatial heterogeneity of host response to SARS-CoV-2 pulmonary infection. Nature Communications, 11(1). https://doi.org/10.1038/s41467-020-20139-7

Diamond, M. S., & Farzan, M. (2013). The broad-spectrum antiviral functions of IFIT and IFITM proteins. In Nature Reviews Immunology (Vol. 13, Issue 1, pp. 46–57). Nat Rev Immunol. https://doi.org/10.1038/nri3344

Diamond, M. S., & Harris, E. (2001). Interferon inhibits dengue virus infection by preventing translation of viral RNA through a PKR-independent mechanism. Virology, 289(2), 297–311. https://doi.org/10.1006/viro.2001.1114

Dionne, K. R., Galvin, J. M., Schittone, S. A., Clarke, P., & Tyler, K. L. (2011). Type I interferon signaling limits reoviral tropism within the brain and prevents lethal systemic infection. Journal of NeuroVirology, 17(4), 314–326. https://doi.org/10.1007/s13365-011-0038-1

Felgenhauer, U., Schoen, A., Gad, H. H., Hartmann, R., Schaubmar, A. R., Failing, K., Drosten, C., & Weber, F. (2020). Inhibition of SARS–CoV-2 by type I and type III interferons. Journal of Biological Chemistry, 295(41), 13958–13964. https://doi.org/10.1074/jbc.AC120.013788

Guan, W., Ni, Z., Hu, Y., Liang, W., Ou, C., He, J., Liu, L., Shan, H., Lei, C., Hui, D. S. C., Du, B., Li, L., Zeng, G., Yuen, K.-Y., Chen, R., Tang, C., Wang, T., Chen, P., Xiang, J., … Zhong, N. (2020). Clinical Characteristics of Coronavirus Disease 2019 in China. New England Journal of Medicine, 382(18), 1708–1720. https://doi.org/10.1056/NEJMoa2002032

Haake, C., Cook, S., Pusterla, N., & Murphy, B. (2020). Coronavirus Infections in Companion Animals: Virology, Epidemiology, Clinical and Pathologic Features. Viruses, 12(9). https://doi.org/10.3390/v12091023

Huang, C., Wang, Y., Li, X., Ren, L., Zhao, J., Hu, Y., Zhang, L., Fan, G., Xu, J., Gu, X., Cheng, Z., Yu, T., Xia, J., Wei, Y., Wu, W., Xie, X., Yin, W., Li, H., Liu, M., … Cao, B. (2020). Clinical features of patients infected with 2019 novel coronavirus in Wuhan, China. The Lancet, 395(10223), 497–506. https://doi.org/10.1016/S0140-6736(20)30183-5

Hung, I. F. N., Lung, K. C., Tso, E. Y. K., Liu, R., Chung, T. W. H., Chu, M. Y., Ng, Y. Y., Lo, J., Chan, J., Tam, A. R., Shum, H. P., Chan, V., Wu, A. K. L., Sin, K. M., Leung, W. S., Law, W. L., Lung, D. C., Sin, S., Yeung, P., … Yuen, K. Y. (2020). Triple combination of interferon beta-1b, lopinavir–ritonavir, and ribavirin in the treatment of patients admitted to hospital with COVID-19: an open-label, randomised, phase 2 trial. The Lancet, 395(10238), 1695–1704. https://doi.org/10.1016/S0140-6736(20)31042-4

Israelow, B., Song, E., Mao, T., Lu, P., Meir, A., Liu, F., Alfajaro, M. M., Wei, J., Dong, H., Homer, R. J., Ring, A., Wilen, C. B., & Iwasaki, A. (2020). Mouse model of SARS-CoV-2 reveals inflammatory role of type i interferon signaling. Journal of Experimental Medicine, 217(12). https://doi.org/10.1084/JEM.20201241

Jilg, N., Lin, W., Hong, J., Schaefer, E. A., Wolski, D., Meixong, J., Goto, K., Brisac, C., Chusri, P., Fusco, D. N., Chevaliez, S., Luther, J., Kumthip, K., Urban, T. J., Peng, L. F., Lauer, G. M., & Chung, R. T. (2014). Kinetic differences in the induction of interferon stimulated genes by interferon-α and interleukin 28B are altered by infection with hepatitis C virus. Hepatology, 59(4), 1250–1261. https://doi.org/10.1002/hep.26653

Katsura, H., Sontake, V., Tata, A., Kobayashi, Y., Edwards, C. E., Heaton, B. E., Konkimalla, A., Asakura, T., Mikami, Y., Fritch, E. J., Lee, P. J., Heaton, N. S., Boucher, R. C., Randell, S. H., Baric, R. S., & Tata, P. R. (2020). Human Lung Stem Cell-Based Alveolospheres Provide Insights into SARS-CoV-2-Mediated Interferon Responses and Pneumocyte Dysfunction. Cell Stem Cell, 27(6), 890–904.e8. https://doi.org/10.1016/j.stem.2020.10.005

Klinkhammer, J., Schnepf, D., Ye, L., Schwaderlapp, M., Gad, H. H., Hartmann, R., Garcin, D., Mahlakõiv, T., & Staeheli, P. (2018). IFN-λ prevents influenza virus spread from the upper airways to the lungs and limits virus transmission. ELife, 7. https://doi.org/10.7554/eLife.33354

Kohli, A., Zhang, X., Yang, J., Russell, R. S., Donnelly, R. P., Sheikh, F., Sherman, A., Young, H., Imamichi, T., Lempicki, R. A., Masur, H., & Kottilil, S. (2012). Distinct and overlapping genomic profiles and antiviral effects of Interferon-λ and -α On HCV-infected and noninfected hepatoma cells. Journal of Viral Hepatitis, 19(12), 843–853. https://doi.org/10.1111/j.1365-2893.2012.01610.x

Lamers, M. M., Beumer, J., Vaart, J. Van Der, Knoops, K., Puschhof, J., Breugem, T. I., Ravelli, R. B. G., Schayck, J. P. Van, Mykytyn, A. Z., Duimel, H. Q., Donselaar, E. Van, Riesebosch, S., Kuijpers, H. J. H., Schipper, D., Wetering, W. J. V. De, Graaf, M. De, Koopmans, M., Cuppen, E., Peters, P. J., … Clevers, H. (2020). SARS-CoV-2 productively infects human gut enterocytes. Science, 369(6499), 50–54. https://doi.org/10.1126/science.abc1669

Lin, J. Da, Feng, N., Sen, A., Balan, M., Tseng, H. C., McElrath, C., Smirnov, S. V., Peng, J., Yasukawa, L. L., Durbin, R. K., Durbin, J. E., Greenberg, H. B., & Kotenko, S. V. (2016). Distinct Roles of Type I and Type III Interferons in Intestinal Immunity to Homologous and Heterologous Rotavirus Infections. PLoS Pathogens, 12(4). https://doi.org/10.1371/journal.ppat.1005600

Lin, L., Jiang, X., Zhang, Z., Huang, S., Zhang, Z., Fang, Z., Gu, Z., Gao, L., Shi, H., Mai, L., Liu, Y., Lin, X., Lai, R., Yan, Z., Li, X., & Shan, H. (2020). Gastrointestinal symptoms of 95 cases with SARS-CoV-2 infection. Gut, 69(6), 997–1001. https://doi.org/10.1136/gutjnl-2020-321013

Lokugamage, K. G., Hage, A., de Vries, M., Valero-Jimenez, A. M., Schindewolf, C., Dittmann, M., Rajsbaum, R., & Menachery, V. D. (2020). Type I interferon susceptibility distinguishes SARS-CoV-2 from SARS-CoV. In bioRxiv. bioRxiv. https://doi.org/10.1101/2020.03.07.982264

Lu, H., Stratton, C. W., & Tang, Y. W. (2020). Outbreak of pneumonia of unknown etiology in Wuhan, China: The mystery and the miracle. In Journal of Medical Virology (Vol. 92, Issue 4, pp. 401–402). John Wiley and Sons Inc. https://doi.org/10.1002/jmv.25678

Luker, G. D., Prior, J. L., Song, J., Pica, C. M., & Leib, D. A. (2003). Bioluminescence Imaging Reveals Systemic Dissemination of Herpes Simplex Virus Type 1 in the Absence of Interferon Receptors. Journal of Virology, 77(20), 11082–11093. https://doi.org/10.1128/jvi.77.20.11082-11093.2003

Mahlakõiv, T., Hernandez, P., Gronke, K., Diefenbach, A., & Staeheli, P. (2015). Leukocyte-Derived IFN-α/β and Epithelial IFN-λ Constitute a Compartmentalized Mucosal Defense System that Restricts Enteric Virus Infections. PLoS Pathogens, 11(4). https://doi.org/10.1371/journal.ppat.1004782

Marcello, T., Grakoui, A., Barba-Spaeth, G., Machlin, E. S., Kotenko, S. V., Macdonald, M. R., & Rice, C. M. (2006). Interferons α and λ Inhibit Hepatitis C Virus Replication With Distinct Signal Transduction and Gene Regulation Kinetics. Gastroenterology, 131(6), 1887–1898. https://doi.org/10.1053/j.gastro.2006.09.052

Meager, A., Visvalingam, K., Dilger, P., Bryan, D., & Wadhwa, M. (2005). Biological activity of interleukins-28 and -29: Comparison with type I interferons. Cytokine, 31(2), 109–118. https://doi.org/10.1016/j.cyto.2005.04.003

Meffre, E., & Iwasaki, A. (2020). Interferon deficiency can lead to severe COVID. In Nature (Vol. 587, Issue 7834, pp. 374–376). NLM (Medline). https://doi.org/10.1038/d41586-020-03070-1

Munis, A. M., Bentley, E. M., & Takeuchi, Y. (2020). A tool with many applications: vesicular stomatitis virus in research and medicine. In Expert Opinion on Biological Therapy (Vol. 20, Issue 10, pp. 1187–1201). Taylor and Francis Ltd. https://doi.org/10.1080/14712598.2020.1787981

Nice, T. J., Baldridge, M. T., McCune, B. T., Norman, J. M., Lazear, H. M., Artyomov, M., Diamond, M. S., & Virgin, H. W. (2015). Interferon-λ cures persistent murine norovirus infection in the absence of adaptive immunity. Science, 347(6219), 269–273. https://doi.org/10.1126/science.1258100

Park, A., & Iwasaki, A. (2020). Type I and Type III Interferons – Induction, Signaling, Evasion, and Application to Combat COVID-19. In Cell Host and Microbe (Vol. 27, Issue 6, pp. 870–878). Cell Press. https://doi.org/10.1016/j.chom.2020.05.008

Pervolaraki, K., Rastgou Talemi, S., Albrecht, D., Bormann, F., Bamford, C., Mendoza, J. L., Garcia, K. C., McLauchlan, J., Höfer, T., Stanifer, M. L., & Boulant, S. (2018). Differential induction of interferon stimulated genes between type I and type III interferons is independent of interferon receptor abundance. PLoS Pathogens, 14(11). https://doi.org/10.1371/journal.ppat.1007420

Pervolaraki, K., Stanifer, M. L., Münchau, S., Renn, L. A., Albrecht, D., Kurzhals, S., Senís, E., Grimm, D., Schröder-Braunstein, J., Rabin, R. L., & Boulant, S. (2017). Type I and type III interferons display different dependency on mitogen-activated protein kinases to mount an antiviral state in the human gut. Frontiers in Immunology, 8(APR). https://doi.org/10.3389/fimmu.2017.00459

Pott, J., Mahlakõiv, T., Mordstein, M., Duerr, C. U., Michiels, T., Stockinger, S., Staeheli, P., & Hornef, M. W. (2011). IFN-λ determines the intestinal epithelial antiviral host defense. Proceedings of the National Academy of Sciences of the United States of America, 108(19), 7944–7949. https://doi.org/10.1073/pnas.1100552108

Pott, J., & Stockinger, S. (2017). Type I and III interferon in the gut: Tight balance between host protection and immunopathology. In Frontiers in Immunology (Vol. 8, Issue MAR). Frontiers Research Foundation. https://doi.org/10.3389/fimmu.2017.00258

Puelles, V. G., Lütgehetmann, M., Lindenmeyer, M. T., Sperhake, J. P., Wong, M. N., Allweiss, L., Chilla, S., Heinemann, A., Wanner, N., Liu, S., Braun, F., Lu, S., Pfefferle, S., Schröder, A. S., Edler, C., Gross, O., Glatzel, M., Wichmann, D., Wiech, T., … Huber, T. B. (2020). Multiorgan and Renal Tropism of SARS-CoV-2. New England Journal of Medicine, 383(6), 590–592. https://doi.org/10.1056/nejmc2011400

Rebendenne, A., Chaves Valadão, A. L., Tauziet, M., Maarifi, G., Bonaventure, B., McKellar, J., Planès, R., Nisole, S., Arnaud-Arnould, M., Moncorgé, O., & Goujon, C. (2021). SARS-CoV-2 Triggers an MDA-5-Dependent Interferon Response Which Is Unable To Control Replication in Lung Epithelial Cells. Journal of Virology, 95(8). https://doi.org/10.1128/jvi.02415-20

Remmelink, M., De Mendonça, R., D’Haene, N., De Clercq, S., Verocq, C., Lebrun, L., Lavis, P., Racu, M. L., Trépant, A. L., Maris, C., Rorive, S., Goffard, J. C., De Witte, O., Peluso, L., Vincent, J. L., Decaestecker, C., Taccone, F. S., & Salmon, I. (2020). Unspecific post-mortem findings despite multiorgan viral spread in COVID-19 patients. Critical Care, 24(1). https://doi.org/10.1186/s13054-020-03218-5

Sadler, A. J., & Williams, B. R. G. (2008). Interferon-inducible antiviral effectors. In Nature Reviews Immunology (Vol. 8, Issue 7, pp. 559–568). Nature Publishing Group. https://doi.org/10.1038/nri2314

Sallard, E., Lescure, F. X., Yazdanpanah, Y., Mentre, F., & Peiffer-Smadja, N. (2020). Type 1 interferons as a potential treatment against COVID-19. Antiviral Research, 178, 104791. https://doi.org/10.1016/j.antiviral.2020.104791

Schaller, T., Hirschbühl, K., Burkhardt, K., Braun, G., Trepel, M., Märkl, B., & Claus, R. (2020). Postmortem Examination of Patients with COVID-19. In JAMA - Journal of the American Medical Association (Vol. 323, Issue 24, pp. 2518–2520). American Medical Association. https://doi.org/10.1001/jama.2020.8907

Selvakumar, T. A., Bhushal, S., Kalinke, U., Wirth, D., Hauser, H., Köster, M., & Hornef, M. W. (2017). Identification of a predominantly interferon-λ-induced transcriptional profile in murine intestinal epithelial cells. Frontiers in Immunology, 8(OCT). https://doi.org/10.3389/fimmu.2017.01302

Sheppard, P., Kindsvogel, W., Xu, W., Henderson, K., Schlutsmeyer, S., Whitmore, T. E., Kuestner, R., Garrigues, U., Birks, C., Roraback, J., Ostrander, C., Dong, D., Shin, J., Presnell, S., Fox, B., Haldeman, B., Cooper, E., Taft, D., Gilbert, T., … Klucher, K. M. (2003). IL-28, IL-29 and their class II cytokine receptor IL-28R. In Nature Immunology (Vol. 4, Issue 1, pp. 63–68). Nat Immunol. https://doi.org/10.1038/ni873

Shuai, H., Chu, H., Hou, Y., Yang, D., Wang, Y., Hu, B., Huang, X., Zhang, X., Chai, Y., Cai, J. P., Chan, J. F. W., & Yuen, K. Y. (2020). Differential immune activation profile of SARS-CoV-2 and SARS-CoV infection in human lung and intestinal cells: Implications for treatment with IFN-β and IFN inducer. Journal of Infection, 81(4), e1–e10. https://doi.org/10.1016/j.jinf.2020.07.016

Stanifer, M. L., Guo, C., Doldan, P., & Boulant, S. (2020). Importance of Type I and III Interferons at Respiratory and Intestinal Barrier Surfaces. In Frontiers in Immunology (Vol. 11, p. 1). Frontiers Media S.A. https://doi.org/10.3389/fimmu.2020.608645

Stanifer, M. L., Kee, C., Cortese, M., Alexandrov, T., Bartenschlager, R., Boulant, S., Zumaran, C. M., Triana, S., Mukenhirn, M., & Kraeusslich, H.-G. (2020). Critical Role of Type III Interferon in Controlling SARS-CoV-2 Infection in Human Intestinal Epithelial Cells ll Critical Role of Type III Interferon in Controlling SARS-CoV-2 Infection in Human Intestinal Epithelial Cells. CellReports, 32, 107863. https://doi.org/10.1016/j.celrep.2020.107863

Stanifer, M. L., Pervolaraki, K., & Boulant, S. (2019). Differential regulation of type I and type III interferon signaling. In International Journal of Molecular Sciences (Vol. 20, Issue 6). MDPI AG. https://doi.org/10.3390/ijms20061445

Tavazzi, G., Pellegrini, C., Maurelli, M., Belliato, M., Sciutti, F., Bottazzi, A., Sepe, P. A., Resasco, T., Camporotondo, R., Bruno, R., Baldanti, F., Paolucci, S., Pelenghi, S., Iotti, G. A., Mojoli, F., & Arbustini, E. (2020). Myocardial localization of coronavirus in COVID-19 cardiogenic shock. European Journal of Heart Failure, 22(5), 911–915. https://doi.org/10.1002/ejhf.1828

Triana, S., Metz-Zumaran, C., Ramirez, C., Kee, C., Doldan, P., Shahraz, M., Schraivogel, D., Gschwind, A. R., Sharma, A. K., Steinmetz, L. M., Herrmann, C., Alexandrov, T., Boulant, S., & Stanifer, M. L. (2021). Single-cell analyses reveal SARS-CoV-2 interference with intrinsic immune response in the human gut. Molecular Systems Biology, 17(4). https://doi.org/10.15252/msb.202110232

Trypsteen, W., Van Cleemput, J., van Snippenberg, W., Gerlo, S., & Vandekerckhove, L. (2020). On the whereabouts of SARS-CoV-2 in the human body: A systematic review. PLoS Pathogens, 16(10). https://doi.org/10.1371/journal.ppat.1009037

Vanderheiden, A., Ralfs, P., Chirkova, T., Upadhyay, A. A., Zimmerman, M. G., Bedoya, S., Aoued, H., Tharp, G. M., Pellegrini, K. L., Manfredi, C., Sorscher, E., Mainou, B., Lobby, J. L., Kohlmeier, J. E., Lowen, A. C., Shi, P.-Y., Menachery, V. D., Anderson, L. J., Grakoui, A., … Suthar, M. S. (2020). Type I and Type III Interferons Restrict SARS-CoV-2 Infection of Human Airway Epithelial Cultures. Journal of Virology, 94(19). https://doi.org/10.1128/jvi.00985-20

Vladimer, G. I., Górna, M. W., & Superti-Furga, G. (2014). IFITs: Emerging roles as key anti-viral proteins. In Frontiers in Immunology (Vol. 5, Issue MAR). Frontiers Research Foundation. https://doi.org/10.3389/fimmu.2014.00094

Voigt, E. A., & Yin, J. (2015). Kinetic Differences and Synergistic Antiviral Effects Between Type I and Type III Interferon Signaling Indicate Pathway Independence. Journal of Interferon and Cytokine Research, 35(9), 734–747. https://doi.org/10.1089/jir.2015.0008

Weber, E., Finsterbusch, K., Lindquist, R., Nair, S., Lienenklaus, S., Gekara, N. O., Janik, D., Weiss, S., Kalinke, U., Overby, A. K., & Kroger, A. (2014). Type I Interferon Protects Mice from Fatal Neurotropic Infection with Langat Virus by Systemic and Local Antiviral Responses. Journal of Virology, 88(21), 12202–12212. https://doi.org/10.1128/jvi.01215-14

Wölfel, R., Corman, V. M., Guggemos, W., Seilmaier, M., Zange, S., Müller, M. A., Niemeyer, D., Jones, T. C., Vollmar, P., Rothe, C., Hoelscher, M., Bleicker, T., Brünink, S., Schneider, J., Ehmann, R., Zwirglmaier, K., Drosten, C., & Wendtner, C. (2020). Virological assessment of hospitalized patients with COVID-2019. Nature, 581(7809), 465–469. https://doi.org/10.1038/s41586-020-2196-x

Wu, N., Nguyen, X. N., Wang, L., Appourchaux, R., Zhang, C., Panthu, B., Gruffat, H., Journo, C., Alais, S., Qin, J., Zhang, N., Tartour, K., Catez, F., Mahieux, R., Ohlmann, T., Liu, M., Du, B., & Cimarelli, A. (2019). The interferon stimulated gene 20 protein (ISG20) is an innate defense antiviral factor that discriminates self versus non-self translation. PLoS Pathogens, 15(10). https://doi.org/10.1371/journal.ppat.1008093

Wu, Y., Guo, C., Tang, L., Hong, Z., Zhou, J., Dong, X., Yin, H., Xiao, Q., Tang, Y., Qu, X., Kuang, L., Fang, X., Mishra, N., Lu, J., Shan, H., Jiang, G., & Huang, X. (2020). Prolonged presence of SARS-CoV-2 viral RNA in faecal samples. In The Lancet Gastroenterology and Hepatology (Vol. 5, Issue 5, pp. 434–435). Elsevier Ltd. https://doi.org/10.1016/S2468-1253(20)30083-2

Xiao, F., Tang, M., Zheng, X., Liu, Y., Li, X., & Shan, H. (2020). Evidence for Gastrointestinal Infection of SARS-CoV-2. Gastroenterology, 158(6), 1831–1833.e3. https://doi.org/10.1053/j.gastro.2020.02.055

Xing, Y. H., Ni, W., Wu, Q., Li, W. J., Li, G. J., Wang, W. Di, Tong, J. N., Song, X. F., Wing-Kin Wong, G., & Xing, Q. S. (2020). Prolonged viral shedding in feces of pediatric patients with coronavirus disease 2019. In Journal of Microbiology, Immunology and Infection (Vol. 53, Issue 3, pp. 473–480). Elsevier Ltd. https://doi.org/10.1016/j.jmii.2020.03.021

Xu, Y., Li, X., Zhu, B., Liang, H., Fang, C., Gong, Y., Guo, Q., Sun, X., Zhao, D., Shen, J., Zhang, H., Liu, H., Xia, H., Tang, J., Zhang, K., & Gong, S. (2020). Characteristics of pediatric SARS-CoV-2 infection and potential evidence for persistent fecal viral shedding. Nature Medicine, 26(4), 502–505. https://doi.org/10.1038/s41591-020-0817-4

Zang, R., Castro, M. F. G., McCune, B. T., Zeng, Q., Rothlauf, P. W., Sonnek, N. M., Liu, Z., Brulois, K. F., Wang, X., Greenberg, H. B., Diamond, M. S., Ciorba, M. A., Whelan, S. P. J., & Ding, S. (2020). TMPRSS2 and TMPRSS4 promote SARS-CoV-2 infection of human small intestinal enterocytes. Science Immunology, 5(47). https://doi.org/10.1126/sciimmunol.abc3582

Zhou, Z., Hamming, O. J., Ank, N., Paludan, S. R., Nielsen, A. L., & Hartmann, R. (2007). Type III Interferon (IFN) Induces a Type I IFN-Like Response in a Restricted Subset of Cells through Signaling Pathways Involving both the Jak-STAT Pathway and the Mitogen-Activated Protein Kinases. Journal of Virology, 81(14), 7749–7758. https://doi.org/10.1128/jvi.02438-06

Zhu, N., Zhang, D., Wang, W., Li, X., Yang, B., Song, J., Zhao, X., Huang, B., Shi, W., Lu, R., Niu, P., Zhan, F., Ma, X., Wang, D., Xu, W., Wu, G., Gao, G. F., & Tan, W. (2020). A Novel Coronavirus from Patients with Pneumonia in China, 2019. New England Journal of Medicine, 382(8), 727–733. https://doi.org/10.1056/nejmoa2001017

